# NSD2 maintains lineage plasticity and castration-resistance in neuroendocrine prostate cancer

**DOI:** 10.1101/2023.07.18.549585

**Authors:** Jia J. Li, Alessandro Vasciaveo, Dimitrios Karagiannis, Zhen Sun, Xiao Chen, Fabio Socciarelli, Ziv Frankenstein, Min Zou, Tania Pannellini, Yu Chen, Kevin Gardner, Brian D. Robinson, Johann de Bono, Cory Abate-Shen, Mark A. Rubin, Massimo Loda, Charles L. Sawyers, Andrea Califano, Chao Lu, Michael M. Shen

**Author notes:** Authors made equal contributions (Jia J. Li, Alessandro Vasciaveo). Addresses for correspondence (C.L.); (M.M.S.).

## Abstract

The clinical use of potent androgen receptor (AR) inhibitors has promoted the emergence of novel subtypes of metastatic castration-resistant prostate cancer (mCRPC), including neuroendocrine prostate cancer (CRPC-NE), which is highly aggressive and lethal^1^. These mCRPC subtypes display increased lineage plasticity and often lack AR expression^2–5^. Here we show that neuroendocrine differentiation and castration-resistance in CRPC-NE are maintained by the activity of Nuclear Receptor Binding SET Domain Protein 2 (NSD2)^6^, which catalyzes histone H3 lysine 36 dimethylation (H3K36me2). We find that organoid lines established from genetically-engineered mice^7^ recapitulate key features of human CRPC-NE, and can display transdifferentiation to neuroendocrine states in culture. CRPC-NE organoids express elevated levels of NSD2 and H3K36me2 marks, but relatively low levels of H3K27me3, consistent with antagonism of EZH2 activity by H3K36me2. Human CRPC-NE but not primary NEPC tumors expresses high levels of NSD2, consistent with a key role for NSD2 in lineage plasticity, and high NSD2 expression in mCRPC correlates with poor survival outcomes. Notably, CRISPR/Cas9 targeting of *NSD2* or expression of a dominant-negative oncohistone H3.3K36M mutant results in loss of neuroendocrine phenotypes and restores responsiveness to the AR inhibitor enzalutamide in mouse and human CRPC-NE organoids and grafts. Our findings indicate that NSD2 inhibition can reverse lineage plasticity and castration-resistance, and provide a potential new therapeutic target for CRPC-NE.

## Introduction

Lineage plasticity has emerged as a central mechanism that drives cancer progression and resistance to therapy, and is now considered a hallmark of cancer^8^. Tumor plasticity represents a formidable challenge for managing cancer care since it contributes to intra-tumor heterogeneity, promotes metastatic dissemination, and enables evasion of targeted therapy. In principle, tumor plasticity can be driven by a range of molecular mechanisms in response to cell-intrinsic changes, such as the acquisition of new mutations or epigenetic modifications, or extrinsic changes including alterations in the microenvironment or treatment regimens. Epigenetic mechanisms that drive lineage plasticity are particularly interesting since they may be more readily reversible and amenable to therapeutic interventions.

In advanced prostate cancer, progression to neuroendocrine castration-resistant prostate cancer (CRPC-NE) occurs through a lineage switch associated with transdifferentiation from luminal adenocarcinoma to neuroendocrine states^2, 7, 9^. Although potent androgen receptor (AR) inhibitors such as abiraterone and enzalutamide have been highly effective for treatment of hormone-sensitive prostate cancer (HSPC), most tumors inevitably develop resistance. Such castration-resistant prostate cancers (CRPC) often retain AR expression and adenocarcinoma histology (CRPC-adeno)^1, 10^. However, metastatic CRPC (mCRPC) can display a wide range of other tumor phenotypes due to lineage plasticity. Notably, CRPC-NE typically lacks AR expression and instead expresses neuroendocrine (NE) markers such as synaptophysin and chromogranin A^4, 5, 11^; occasionally, it can also occur in an amphicrine form that expresses both AR and NE markers^12^. Yet another subtype that has been distinguished in mCRPC is double-negative prostate cancer (DNPC), which lacks both AR and NE marker expression^13, 14^. Importantly, the classical form of neuroendocrine prostate cancer that arises *de novo* in primary tumors (primary NEPC) in the absence of androgen-deprivation is relatively rare (less than 0.1%^15^), whereas CRPC-NE occurs in at least 10-25% of mCRPC^2, 3, 16–18^.

There is considerable evidence that the lineage plasticity in CRPC-NE is mediated by epigenetic reprogramming^2, 3, 5, 11, 19–21^. Studies from multiple laboratories have shown that CRPC-NE is driven by loss-of-function of the tumor suppressors *TP53, RB1,* and *PTEN,* which facilitate epigenetic reprogramming and lineage plasticity^2, 4, 22, 23^. In particular, mCRPC expresses high levels of EZH2^24^, the enzymatic subunit of the Polycomb Repressive Complex 2 (PRC2), which mediates tri-methylation of histone H3 lysine 27 (H3K27me3) at the enhancers and promoters of downstream target genes. Several studies have reported that EZH2 promotes neuroendocrine differentiation in CRPC through repression of AR and luminal adenocarcinoma differentiation programs^25–27^.

Despite the significance of CRPC-NE, there has been a relative lack of useful model systems that accurately recapitulate the cell state transitions in neuroendocrine transdifferentiation and enable molecular analyses of the causal mechanisms. In our study, we have developed new mouse organoid models to demonstrate that the histone methyltransferase NSD2 is required for maintenance of neuroendocrine differentiation as well as castration-resistance in CRPC-NE. NSD2 catalyzes the formation of H3K36me2^28–30^, a histone post-translational modification associated with active chromatin. H3K36me2 has been reported to antagonize the activity of PRC2^31–34^, and recruit the *de novo* DNA methyltransferase DNMT3A^35–37^, but its role in transcriptional regulation remains only partially understood.

Here we show that lineage plasticity in CRPC-NE is modulated by H3K36me2 marks generated by NSD2. Genetic knock-out or oncohistone-mediated inhibition of NSD2 can revert neuroendocrine differentiation in CRPC-NE organoids and grafts, and remarkably can re-establish response to the AR inhibitor enzalutamide. Thus, NSD2 targeting may represent an effective therapeutic strategy for CRPC-NE.

## Results

### Mouse organoid lines that recapitulate heterogeneity of human CRPC

To study lineage plasticity in CRPC, we have established tumor organoid lines from *Nkx3.1^CreERT2/+^; Pten^flox/flox^; TrpP53^flox/flox^; Rosa26-EYFP (NPp53)* mice. In this model, tamoxifen induction of the *Nkx3.1^CreERT^*^2^ allele^38^ in adult mice results in specific deletion of the *Pten* and *Trp53* tumor suppressor genes in distal luminal epithelial cells of the prostate. These *NPp53* mice develop CRPC that can undergo transdifferentiation of luminal adenocarcinoma cells to CRPC-NE, as shown by lineage-tracing of the *Rosa26-EYFP* reporter^7^. We established tumor organoid lines from 21 independent *NPp53* mice that had been induced by tamoxifen treatment at 10-12 weeks of age and harvested between 7-15 months of age (Supplementary Table 1). At these stages, the *NPp53* tumors resemble highly aggressive CRPC that is insensitive to AR inhibitors such as abiraterone or enzalutamide^7^.

Of these 21 organoid lines, 6 contained cells with neuroendocrine (NE) features as determined by histopathology and marker expression and were denoted NPPO-1 (Nkx3.1, Pten, P53 Organoid line −1) through NPPO-6 (Fig. 1a; Extended Data Fig. 1; Supplementary Table 2); an additional 3 CRPC lines that lacked NE features were denoted NPPO-7 through NPPO-9 (Extended Data Fig. 2a). Apart from NPPO-3, which could not be maintained after the first passage, the other 5 NPPO lines have been stably maintained for over 20 passages in culture. Notably, the NPPO-1 through NPPO-6 lines displayed distinctive and unique phenotypes that recapitulate much of the spectrum of human CRPC and CRPC-NE (Fig. 1a). For example, the NPPO-1 and NPPO-5 organoid lines displayed phenotypic heterogeneity, containing AR-negative cells with a small cell histology that expressed the NE markers Chromogranin A (CHGA) and Synaptophysin (SYP), together with mesenchymal-like cells that were positive for AR and Vimentin (VIM). The NPPO-2 line was relatively homogeneous, with most cells positive for CHGA and SYP and little or no AR; the NPPO-3 line was similar except that CHGA and SYP expression was less uniform. In contrast, the NPPO-4 and NPPO-6 lines co-expressed NE markers together with AR, and could be considered amphicrine, with the NPPO-6 line displaying a large cell histology.

**Fig. 1.**
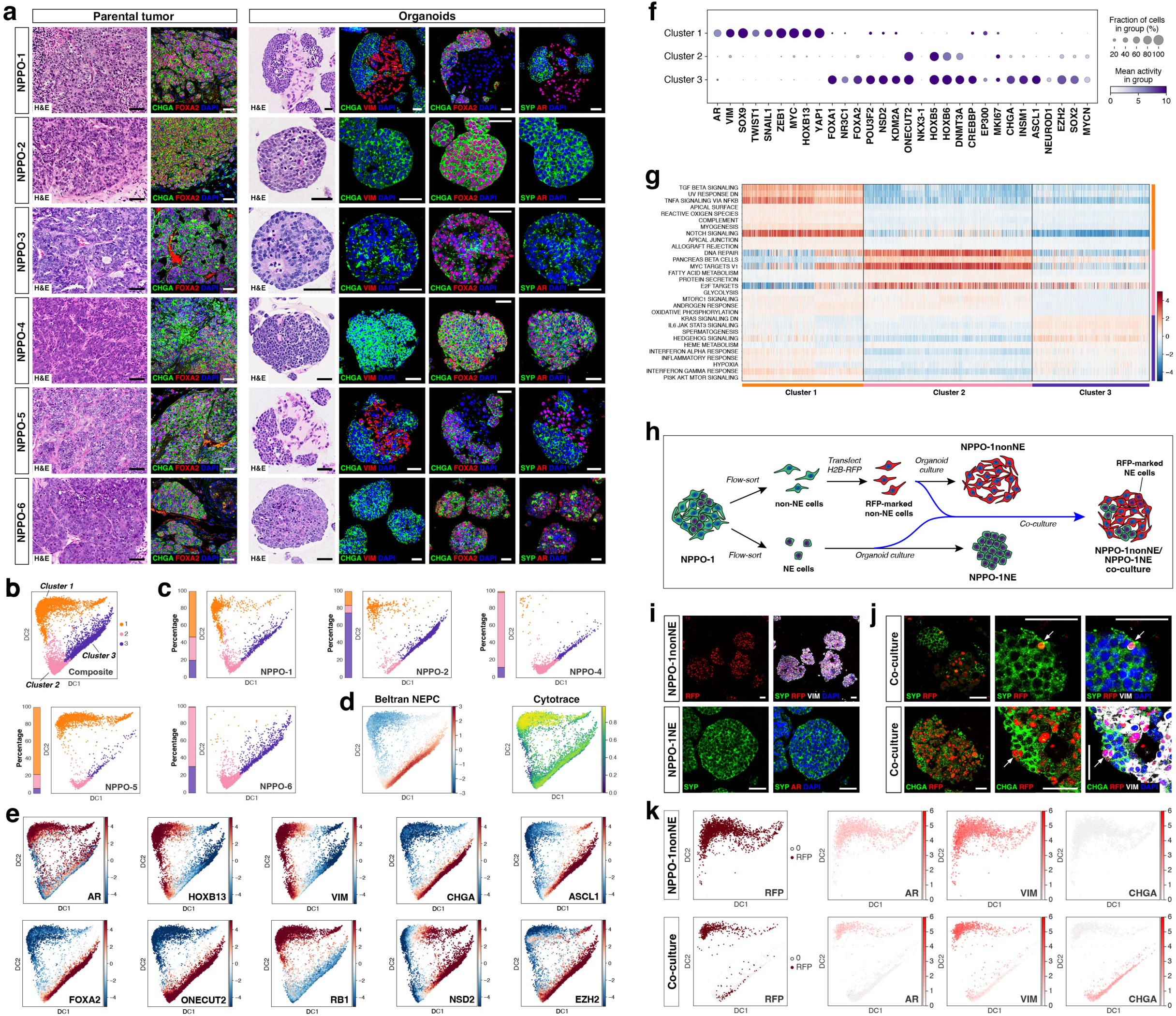
Organoids from *NPp53* mice recapitulate heterogeneity of CRPC-NE. **a,** Hematoxylin and eosin (H&E) and immunofluorescence staining of sections from parental tumors and matched NPPO organoid lines established from *NPp53* mice at passage 2. AR, Androgen receptor; CHGA, Chromogranin A; SYP, Synaptophysin; VIM, Vimentin. **b,** Diffusion Component (DC) projection of scRNA-seq data from composite dataset of all 5 NPPO organoid lines. **c,** DC projection of individual NPPO organoid lines analyzed by VIPER. Proportions of cells in the three clusters are indicated by bars at left of each plot. **d,** *(left)* VIPER-inferred activity for a published NEPC gene signature^23^ and *(right)* CytoTRACE analysis. **e,** Activity profiles inferred by VIPER for the indicated proteins in the composite dataset. **f,** Dot-plot of inferred protein activities in the three clusters. **g,** Pathway analysis using VIPER inferred activities, with the 10 most significantly enriched pathways shown for each cluster. **h,** Schematic depiction of co-culture assay for neuroendocrine transdifferentiation. **i,j,** Immunofluorescence analysis of RFP-expressing NPPO-1nonNE and NPPO-1NE organoids cultured separately (**i**) and together in organoid chimeras (**j**) for 4 passages. Arrows indicate cells co-expressing RFP and VIM together with the neuroendocrine markers CHGA and SYP. **k,** VIPER analysis of scRNA-seq data from NPPO-1nonNE *(top)* and co-cultured organoids *(bottom)* at passage 4; note presence of RFP-positive cells in Clusters 2 and 3 in the co-cultured organoids. Scale bars in **a,i,j** indicate 50 microns.

To assess the heterogeneity and transcriptomic programs of these organoid lines, we performed single-cell RNA-sequencing (scRNA-seq) of early passage NPPO-1, −2, −4, −5, and −6 organoid lines (Supplementary Table 3), followed by protein activity inference by regulatory network analysis with the VIPER algorithm^39^. VIPER provides single-cell measurements of the differential activity of regulatory proteins based on the differential expression of their downstream gene targets (regulon), reverse-engineered using the ARACNe algorithm^40, 41^. These analyses identified three major cell clusters that were present in all 5 organoid lines to varying degrees (Fig. 1b,c). In particular, we observed that Clusters 2 and 3 showed enrichment for a human CRPC-NE signature^23^, whereas Cluster 1 had the highest Cytotrace score, which is a measurement of transcriptional diversity associated with stem/progenitor-like properties^42^ (Fig. 1d). Cluster 1 had high activity of AR and luminal epithelial markers such as HOXB13 and SOX9, as well as mesenchymal markers and regulators of epithelial-mesenchymal transition (EMT), such as VIM, TWIST1, and ZEB1 (Fig. 1e,f; Extended Data Fig. 2b), thereby resembling human CRPC-adeno and also “mesenchymal stem-like prostate cancer” (MSPC)^43^. Cluster 2 lacked activity of AR and displayed low-moderate activity of most NE markers, whereas Cluster 3 resembled CRPC-NE, with high activity of CHGA, ASCL1, and FOXA2; interestingly, a subpopulation of Cluster 3 displayed AR activity, potentially corresponding to AMPC. Finally, analyses of pathway enrichment revealed that Notch signaling was upregulated in Cluster 1 but low in Cluster 3, consistent with its neuroendocrine features (Fig. 1g), and that Cluster 2 displayed strong enrichment for MYC targets as well as E2F targets, consistent with low RB1 activity (Fig. 1e,g).

### Transdifferentiation to neuroendocrine states in organoid culture

The phenotypic stability of the heterogeneous NPPO-1 organoid line suggested that its neuroendocrine (NE) and non-neuroendocrine (non-NE) populations might be maintained by paracrine interactions and/or cell state interconversions. Therefore, we used lineage-tracing to investigate whether non-NE cells could transition to NE states in culture, paralleling the transdifferentiation observed in *NPp53* tumors *in vivo*^7^. Using flow cytometry, we purified separate NE and non-NE populations from NPPO-1 organoids, resulting in isogenic NPPO-1NE and NPPO-1nonNE sublines (Extended Data Figs. 2a, 3a; Methods). We then introduced a H2B-RFP expression cassette by lentiviral infection to mark the NPPO-1nonNE cells with 70% labeling efficiency (Fig. 1h). After 4 passages, we found that NPPO-1NE and NPPO-1nonNE cells cultured separately as organoids were homogeneously neuroendocrine and non-neuroendocrine, respectively (Fig. 1i). In contrast, organoids derived from co-culture of NPPO-1NE and RFP-marked NPPO-1nonNE cells contained rare RFP-expressing cells that gained SYP or CHGA expression, indicating a shift from mesenchymal to NE states (Fig. 1j). VIPER analyses of scRNA-seq data confirmed that the NPPO-1nonNE organoids lacked NE marker expression, whereas the co-cultured organoids contained both nonNE and NE populations, with many RFP-expressing cells in Clusters 2 and 3 (Fig. 1k). These lineage-tracing data indicate that paracrine interactions can promote a cell state transition from an AR-positive mesenchymal state to a NE state in organoid culture.

### NSD2 and H3K36me2 are upregulated in neuroendocrine tumor cells

To investigate potential epigenetic mechanisms that drive neuroendocrine differentiation, we compared the levels of histone modifications in NE cells versus non-NE cells of heterogeneous NPPO-1 organoids by immunofluorescence staining. Although most histone marks examined displayed similar abundance, we observed differential levels for histone H3 lysine 36 dimethylation (H3K36me2), histone H3 lysine 27 acetylation (H3K27ac), and histone H3 lysine 27 trimethylation (H3K27me3); notably, no differences were found in the levels of H3K36me3 (Extended Data Fig. 4a,b). Based on these findings, we focused on H3K36me2 and the histone methyltransferases of the *NSD* family that can catalyze its formation. Consequently, we assessed the relative expression levels of *NSD1, NSD2,* and *NSD3* in human CRPC-adeno and CRPC-NE samples from a published dataset^23^, and found that only *NSD2* was differentially expressed at higher levels in CRPC-NE (Extended Data Fig. 4c).

We next performed Western blotting of 4 organoid lines with NE features and 4 non-NE organoid lines. Notably, we found that H3K36me2 and H3K27ac were upregulated in all NE lines, whereas H3K27me3 displayed the opposite pattern (Fig. 2a). Interestingly, we observed higher levels of both NSD2 and EZH2 in the NE organoid lines versus the non-NE lines, indicating that EZH2 expression does not correlate with the levels of H3K27me3 (Fig. 2b). Thus, EZH2 expression is relatively high in the NE organoid lines, whereas the global levels of H3K27me3 marks are relatively low, consistent with the role of H3K36me2 in antagonizing PRC2 activity^33, 34^.

**Fig. 2.**
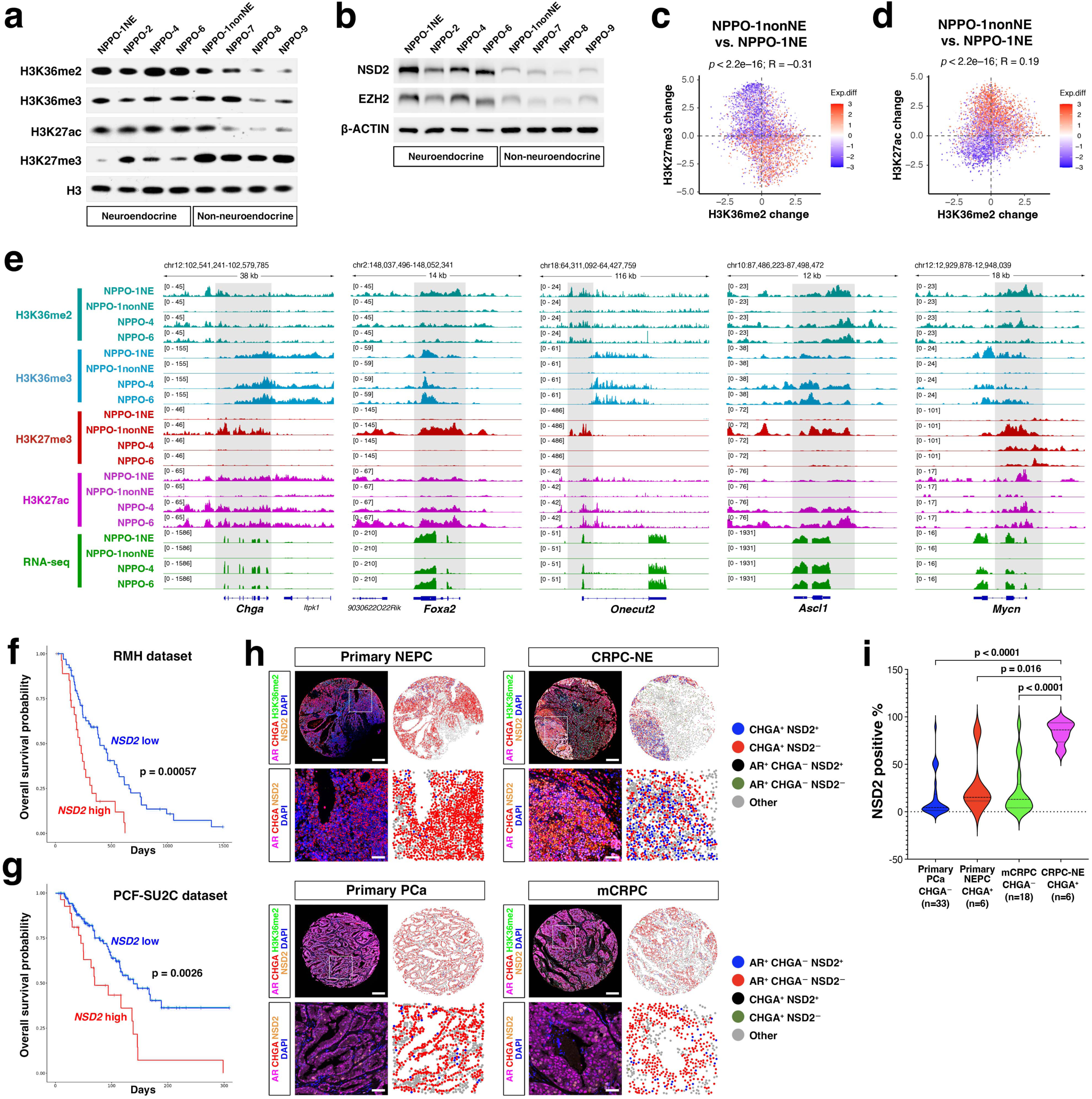
NSD2 is upregulated in CRPC-NE and correlates with poor outcomes. **a,b,** Western blotting for indicated histone marks and NSD2 and EZH2 proteins in neuroendocrine and non-neuroendocrine organoid lines. **c,d,** Correlation of genome-wide differential H3K36me2 and H3K27me3 marks (**c**) or H3K36me2 and H3K27ac marks (**d**) with differential gene expression levels in NPPO-1nonNE versus NPPO-1NE. One-way ANOVA was used to determine differences between groups. **E,** Genome browser view of CUT&Tag signals for H3K36me2, H3K36me3, H3K27ac, and H3K27me3 together with bulk RNA-seq reads at *Chga, Foxa2, Onecut2, Ascl1,* and *Mycn* loci in the indicated organoid lines. **f,g,** Correlation of high NSD2 expression with worse survival outcomes in two independent mCRPC patient cohorts from the Royal Marsden Hospital (RMH; n=28/94) and Prostate Cancer Foundation-Stand Up to Cancer Dream Team (PCF-SU2C; n=27/141). **h,i,** Analysis of prostate cancer tissue microarray. **h,** Five-color overlay images of representative tissue cores and high-power views of four-color images. Spatial plots show the enrichment of NSD2^+^ cells among CHGA^+^ neuroendocrine tumor cells or among AR^+^/CHGA^−^ tumor cells. Scale bars indicate 50 microns. **i,** Violin plot for the percentage of NSD2^+^ cells in CHGA^+^ neuroendocrine tumor cells or AR^+^/CHGA^−^ tumor cells in each patient. Welch’s ANOVA followed by Dunnett’s Multiple Comparison test was used for multiple comparisons among different patient groups.

These findings were confirmed in an independent set of mouse organoid lines with adenocarcinoma and neuroendocrine prostate cancer phenotypes established from *Trp53^flox/flox^*; *Rb1^flox/flox^*; *Pten^flox/flox^* (TKO) mice. In contrast with the NPPO lines, *Trp53*, *Rb1* and *Pten* were deleted in TKO organoids by *ex vivo* infection with Cre-expressing virus and then transplanted *in vivo* (orthotopic or subcutaneously), whereupon we observed prostate adenocarcinomas that transition to neuroendocrine phenotypes. Explants from these grafts cultured as organoids displayed stable adenocarcinoma and neuroendocrine phenotypes (Extended Data Fig. 2c). Notably, Western blotting of TKO organoids showed high levels of NSD2 and H3K36me2 as well as lower levels of H3K27me3 in the neuroendocrine lines relative to the normal and isogenic adenocarcinoma organoid lines (Extended Data Fig. 5a).

Next, we performed genome-wide profiling of histone marks using Cleavage Under Targets & Tagmentation (CUT&Tag)^44^ together with bulk RNA-seq. At the genome-wide level, we observed higher levels of H3K36me2 in neuroendocrine organoid lines were associated with increased levels of target gene expression, together with lower levels of H3K27me3 and increased H3K27ac (Fig. 2c,d; Extended Data Fig. 5b-d). In particular, the H3K36me2 mark was enriched in the putative enhancer (marked by non-promoter H3K27ac peaks) and promoter proximal regions^45^ for actively transcribed NE markers and regulators in the NE organoid lines, including *Chga, Foxa2, Onecut2, Ascl1, Mycn, Insm1,* and *Syp* (Fig. 2e; Extended Data Fig. 5e); we also observed H3K36me3 enrichment in gene body regions of these actively transcribed genes. In contrast, the H3K27me3 mark generally displayed a reciprocal enrichment at these loci in the NPPO-1nonNE line, consistent with its lack of NE marker expression (Fig. 2e; Extended Data Fig. 5e).

To determine the extent and significance of high NSD2 expression in human prostate cancer, we examined its correlation with patient outcomes. Using bulk RNA-seq data from two independent mCRPC cohorts, we found that high NSD2 expression in the Royal Marsden Hospital cohort (n=28/94) as well as the Prostate Cancer Foundation-Stand Up to Cancer cohort (n=27/141) was significantly correlated with poor overall survival (Fig. 2f,g).

Finally, we examined the levels of NSD2 and H3K36me2 in human prostate tumors by multiplexed immunofluorescence on a tissue microarray (TMA). Of the 63 samples on this TMA, 33 were treatment-naïve primary prostate cancer (PCa), 6 were *de novo* primary NEPC, 18 were mCRPC tumors lacking NE features, and 6 were CRPC-NE (Supplementary Table 4). Our analyses revealed that NSD2 was upregulated in all 6 of the CRPC-NE tumors, but notably in only 1 of the 6 primary NEPC samples (Fig. 2h,i). We also observed a similar but less pronounced increase in H3K36me2 levels in the CRPC-NE samples relative to the mCRPC samples (Extended Data Fig. 5f,g). These results are consistent with previous TMA analyses that have demonstrated low NSD2 expression in primary tumors compared to high expression in advanced disease stages^46–48^. However, our analyses also reveal that high levels of NSD2 and H3K36me2 are associated with CRPC-NE and not primary NEPC, suggesting that NSD2 expression in prostate cancer is correlated with lineage plasticity, particularly transdifferentiation towards neuroendocrine states.

### NSD2 inhibition in CRPC-NE reverts neuroendocrine phenotypes

To investigate the functional role of NSD2 in maintaining the neuroendocrine phenotype of NPPO organoids, we performed CRISPR/Cas9-mediated targeting by lentiviral transfection of control (sgCtrl) and targeted (sgNsd2) guides and isolation of infected cells by flow-sorting (Extended Data Fig. 3b). Following CRISPR-mediated knock-out of *Nsd2,* we observed histological alterations and numerous AR-positive cells that were negative for NE markers in the NPPO-1NE and NPPO-2 organoid lines, consistent with conversion of neuroendocrine cells to AR-positive adenocarcinoma cells following NSD2 loss (Fig. 3a,c; Supplementary Table 2); however, no effects were observed in the NPPO-4 and NPPO-6 lines. Therefore, we further investigated the role of H3K36me2 marks using the oncohistone H3.3K36M mutant, which has a dominant-negative effect on NSD family and SETD2 methyltransferases and results in depletion of H3K36me2 and H3K36me3^49–51^. We found that lentiviral expression of H3.3K36M led to lethality of NPPO-1NE and NPPO-2 organoids, but resulted in histological alterations and upregulation of AR in NPPO-6 organoids as well as dramatic reduction of NE markers and AR upregulation in NPPO-4 organoids (Fig. 3e; Extended Data Figs. 3c and 6a; Supplementary Table 2).

**Fig. 3.**
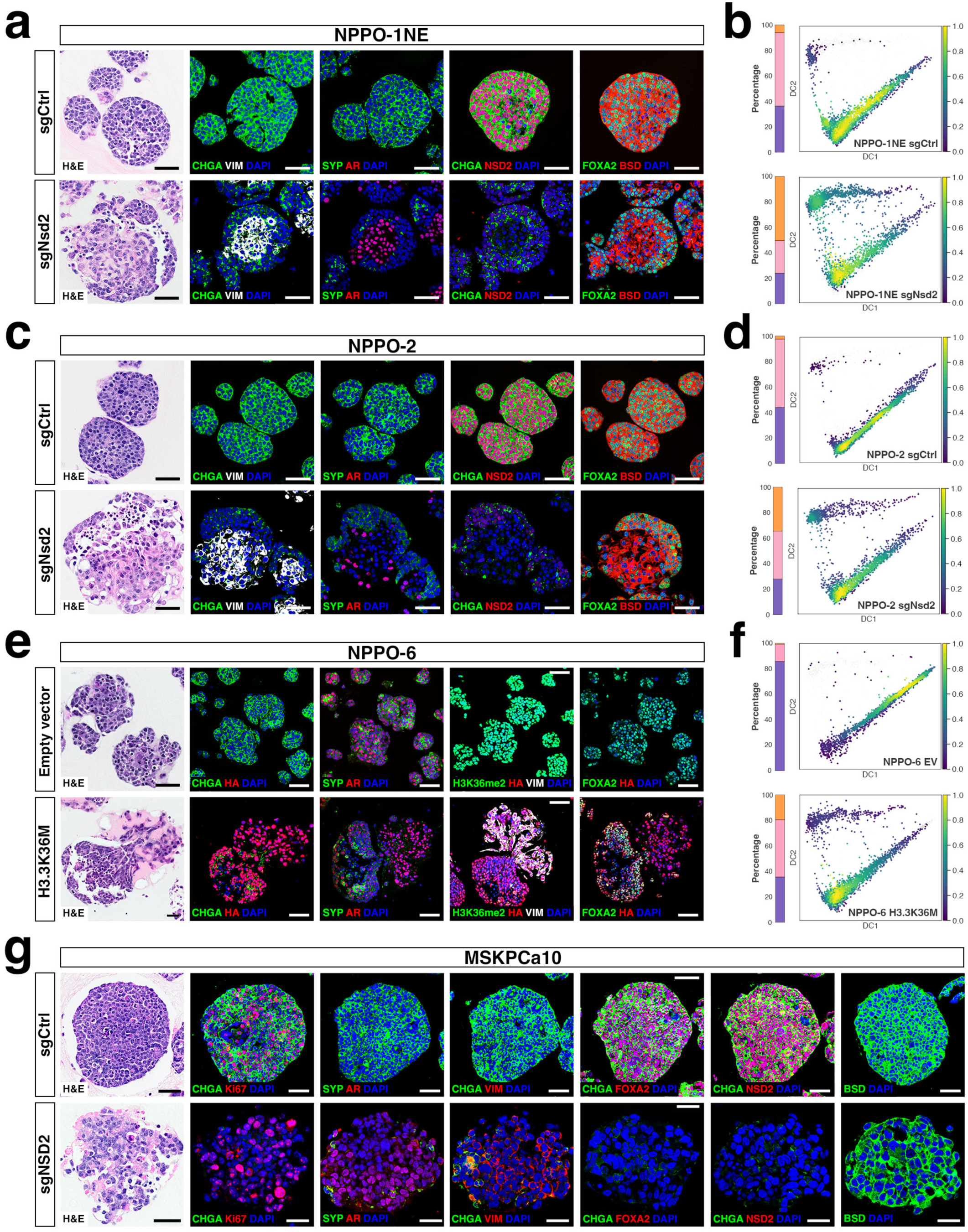
NSD2 inhibition reverts neuroendocrine differentiation. **a,c,e,g,** Hematoxylin-and-eosin (H&E) staining and immunofluorescence staining of sections from NPPO-1NE (**a**), NPPO-2 (**c**), or MSKPCa10 (**g**) organoids following CRISPR-mediated knock-out of *Nsd2* (sgNsd2) or control (sgCtrl), or NPPO-6 organoids (**e**) after transfection of the oncohistone H3.3K36M or control (EV, empty vector). AR, Androgen receptor; BSD, Blasticidin (drug selection marker); CHGA, Chromogranin A; CK8, Cytokeratin 8; HA, Hemagglutinin tag; SYP, Synaptophysin, VIM, Vimentin. **b,d,f,** Density plots for VIPER-analyzed scRNA-seq data from the indicated organoid lines following *Nsd2* knock-out or H3.3K36M expression. Changes in cluster sizes are quantified in vertical bars at left of each plot. Scale bars in panels **a,c,e,g** indicate 50 microns.

To confirm these findings at the single-cell level, we performed multiome single-nuclei RNA-seq/ATAC-seq on the NPPO-1NE and NPPO-2 *Nsd2* targeted organoids and controls, as well as the NPPO-6 H3.3K36M organoids and controls (Supplementary Table 3). VIPER analysis of these snRNA-seq data showed that *Nsd2* targeting or H3.3K36M expression resulted in a substantial population shift from Clusters 2 and 3 toward Cluster 1, consistent with the loss of cells expressing NE markers and gain of AR-positive cells (Fig. 3b,d,f). Western blotting confirmed that *Nsd2* targeting led to loss of NSD2 expression and that NSD2 loss as well as H3.3K36M expression resulted in decreased H3K36me2 and increased H3K27me3 levels (Extended Data Fig. 6b). Similarly, CUT&Tag analyses of the NPPO-1NE, NPPO-2, NPPO-4, and NPPO-6 treatment pairs showed decreased global levels of H3K36me2 and increased H3K27me3 after NSD2 inhibition by knock-out or H3.3K36M expression, with relatively modest effects on H3K36me3 (Extended Data Fig. 6d-h). Finally, we identified differentially active regulatory genes affected by alterations of H3K36me2 histone marks by integration of VIPER-processed snRNA-seq data with bulk CUT&Tag analyses (Extended Data Fig. 7).

To confirm these findings in human CRPC-NE, we utilized the MSKPCa10 organoid line, which is mutant for *TP53* and lacks *Rb1* expression^52^. CRISPR-mediated targeting of *NSD2* resulted in loss of CHGA- and SYP-positive neuroendocrine cells and gain of AR expression, as well as histopathological changes with a decreased nucleus to cytoplasm (N:C) ratio (Fig. 3g). Western blotting showed decreased levels of NSD2 and H3K36me2, as well as increased H3K27me3 levels (Extended Data Fig. 6c), consistent with our findings with mouse NPPO organoids.

### NSD2 inhibition restores enzalutamide response in CRPC-NE

Since *Nsd2* targeting or H3.3K36M expression could restore AR expression in CRPC-NE organoids, we examined whether response to the AR inhibitor enzalutamide would also be enhanced. We found that control NPPO organoids were highly resistant to enzalutamide, whereas *Nsd2* targeting or H3.3K36M expression resulted in significant growth reduction in the presence of enzalutamide, resulting in IC_50_ values of approximately 3 µM or less (Fig. 4a; Extended Data Fig. 8a-f). To confirm these findings *in vivo,* we performed subcutaneous allografting of organoids in immunodeficient NOD/SCID mice, followed by treatment of host mice with enzalutamide or DMSO control after tumors reached approximately 200-250 mm^3^ in size at two weeks after grafting. Compared to controls, enzalutamide treatment significantly reduced the growth of *Nsd2* targeted NPPO-1NE and NPPO-2 grafts as well as H3.3K36M-expressing NPPO-4 and NPPO-6 grafts (Fig. 4b; Extended Data Fig. 9a). Analysis of graft sections by H&E staining and immunofluorescence showed that NSD2 inhibition resulted in loss of NE marker expression, decreased Ki67 expression, and gain of adenocarcinoma phenotypes (Fig. 4c,d; Extended Data Fig. 9b; Supplementary Table 2). Similar results were observed with NSD2 targeting in human MSKPCa10 organoids, which displayed decreased growth after enzalutamide treatment of organoids and xenografts (Fig. 4e-g; Extended Data Figs. 8h,i and 9a; Supplementary Table 2). Taken together, these results indicate that NSD2 inhibition renders CRPC-NE more responsive to enzalutamide.

**Fig. 4.**
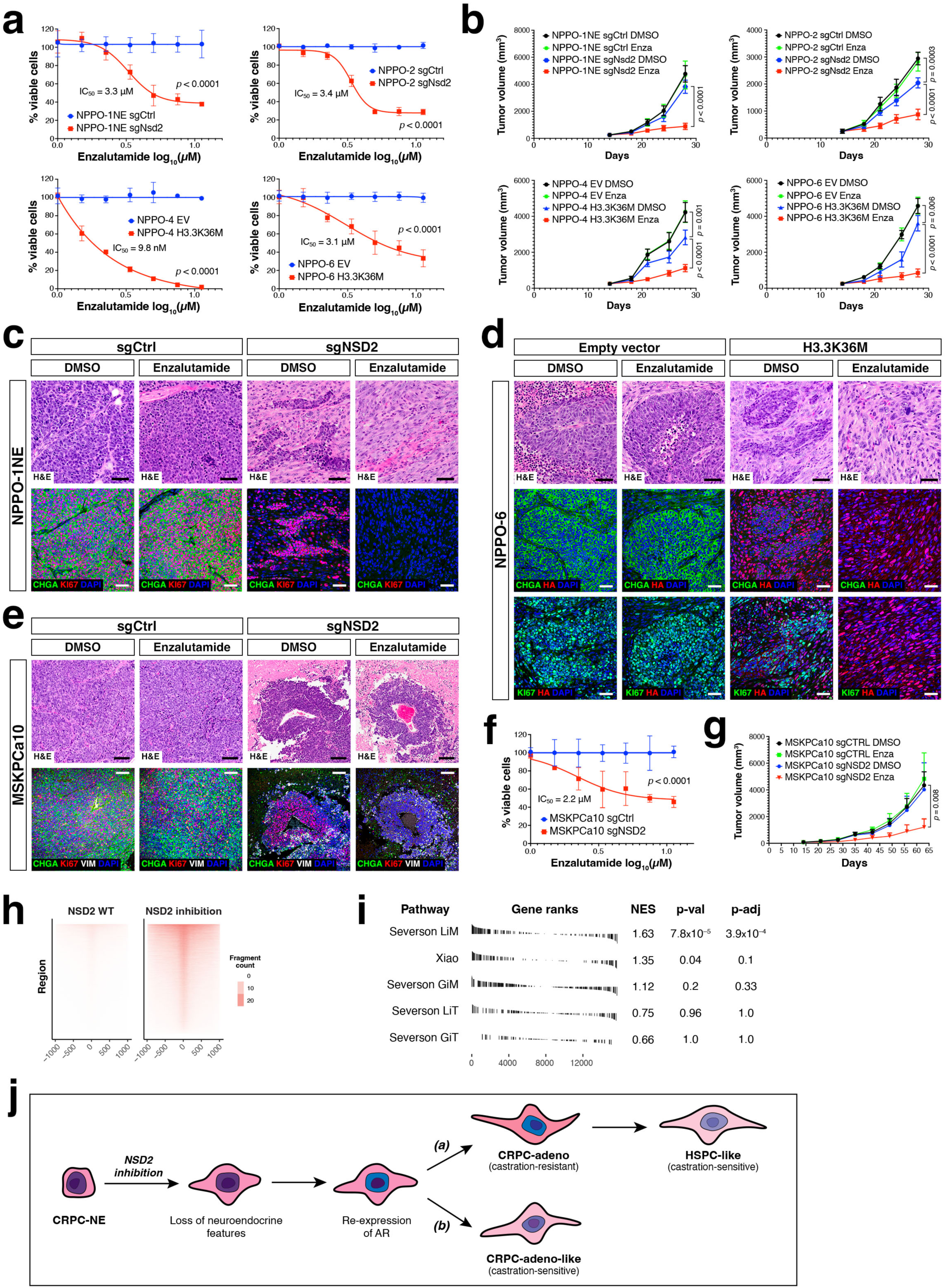
NSD2 inhibition restores enzalutamide response in neuroendocrine organoids and grafts. **a,** Dose-response curves for enzalutamide treatment of *Nsd2* knock-out (sgNsd2) or control (sgCtrl) NPPO-1NE and NPPO-2 organoids and for control (EV, empty vector) or H3.3K36M-transfected NPPO-4 and NPPO-6 organoids. IC_50_ values were calculated from dose-response curves by nonlinear regression (curve fit). Each data point corresponds to three biological replicates; error bars represent one standard deviation. Dose-response curves were compared by two-way ANOVA. **b,** Tumor growth curves for the same lines as in **a**, except treated with enzalutamide or DMSO control *in vivo* starting at day 14 after subcutaneous grafting in NOD/SCID mice at day 0. Each data point corresponds to five biological replicates; error bars represent one standard deviation. Unpaired t-tests (two-tailed P value) were used to compare the means between two groups. **c,d,e,** H&E and immunofluorescence analysis of sections from NPPO-1NE (**c**), NPPO-6 (**d**), and MSKPCa10 grafts (**e**). CHGA, Chromogranin A; HA, Hemagglutinin tag. **f,** Dose-response curve for enzalutamide treatment of *NSD2* knock-out (sgNSD2) or control (sgCtrl) MSKPCa10 organoids. IC_50_ values and statistics were calculated as in panel **a**. **g,** Tumor growth curves for control and *NSD2* knock-out MSKPCa10 subcutaneous xenografts treated with enzalutamide or DMSO control *in vivo*. Each data point corresponds to six biological replicates; error bars represent one standard deviation. Unpaired t-tests (two-tailed P value) were used to compare the means between two groups. **h,** Tornado plots of chromatin accessibility around canonical AR binding sites in AR-expressing cells in scATAC-seq data from combined multiome datasets for *Nsd2* knock-out NPPO-1NE and NPPO-2 and H3.3K36M-transfected NPPO-6 as well as a dataset from untransfected NPPO-1. **i,** Gene set enrichment analysis of signatures for AR target genes, comparing combined control and experimental datasets. Enrichments for the Severson “lost in metastasis” (LiM)^55^ and Xiao^56^ signatures are significant. **j,** Model for loss of neuroendocrine differentiation and castration-resistance after NSD2 inhibition. See text for description.

To investigate the basis for the restoration of enzalutamide sensitivity following NSD2 inhibition, we used our multiome snATAC-seq data from NPPO-1NE, NPPO-2 and NPPO-6 organoids to examine chromatin accessibility at AR target genes in AR-positive cells. We found increased chromatin accessibility at genomic loci around canonical AR binding sites after NSD2 inhibition (Fig. 4h), consistent with re-activation of canonical AR targets. In parallel, we used the snRNA-seq data to examine the expression of published AR target gene signatures^53–56^. We found that a signature of AR target genes that were downregulated in metastases (LiM; lost in metastases^55^) was significantly enriched in AR-positive cells following NSD2 inhibition (Fig. 4i), indicating re-expression of AR targets that are expressed in HSPC but not mCRPC; similar enrichment was observed using an independent AR target gene signature^56^. Given the extensive reprogramming of the AR cistrome toward non-canonical targets during progression to mCRPC^53, 54, 57^, these findings suggest that NSD2 inhibition can shift the AR cistrome towards an HSPC-like state. Taken together, our findings suggest that targeting of NSD2 in CRPC-NE can reverse lineage plasticity, resulting in loss of neuroendocrine phenotypes and response to treatment with androgen receptor inhibitors.

## Discussion

Our findings demonstrate that NSD2 plays a critical role in maintenance of neuroendocrine states and castration-resistance in CRPC-NE. In particular, loss of NSD2 results in reversal of neuroendocrine differentiation and alteration of the lineage plasticity of CRPC-NE. These activities of NSD2 in lineage plasticity are correlated with epigenetic reprogramming and transcriptional activation of key CRPC-NE regulators including FOXA2 and ONECUT2^58, 59^. However, since NSD2 is not broadly upregulated in primary NEPC, NSD2 activity does not appear to be a feature of neuroendocrine states in general, but instead is associated with the lineage plasticity found in CRPC.

Importantly, inhibition of NSD2 in CRPC-NE also results in re-expression of AR, but this NSD2-deficient state is sensitive to the AR inhibitor enzalutamide, unlike CRPC-adeno which is castration-resistant. Thus, we can envision two conceptual models for how NSD2 loss might result in restoration of enzalutamide sensitivity (Fig. 4j). One possibility is NSD2 has a unitary role in modulating lineage plasticity in CRPC, such that NSD2 loss would fully revert CRPC-NE back to an AR-positive cellular state that resembles hormone-sensitive prostate cancer (HSPC). Alternatively, loss of NSD2 might revert CRPC-NE to a state similar to CRPC-adeno that has acquired sensitivity to AR inhibitors, due to an activity of NSD2 in maintaining castration-resistance that is independent of its role in lineage plasticity.

With respect to this second model, NSD2 might facilitate castration-resistance through a protein-protein interaction between NSD2 and AR that is mediated by the HMG domain of NSD2 and thereby alters AR transcriptional activity^60^. Thus, NSD2 loss could affect AR binding properties, consistent with the observed alterations in the AR cistrome following NSD2 inhibition. If this model is correct, NSD2 may have a broader requirement in castration-resistance for at least a subset of non-neuroendocrine mCRPC with higher NSD2 levels, which remains to be evaluated. Furthermore, NSD2 has been previously implicated in promoting metastasis of prostate cancer^46, 61, 62^, and may thereby represent a functional link between lineage plasticity, castration-resistance, and metastasis in mCRPC.

The functions of NSD2 in prostate cancer are consistent with a general role of H3K36me2 marks and NSD histone methyltransferases in promoting lineage plasticity and metastasis in a range of tumor types^63^. Notably, NSD2 is activated by a t(4,14) translocation in a major subtype of multiple myeloma^64^, and its gain-of-function mutations are frequently observed in pediatric acute lymphoblastic leukemia^65^. The related genes NSD1 and NSD3 have also been shown to be key drivers of head and neck cancer and squamous cell lung cancer, respectively^66–68^. Furthermore, NSD2 may be therapeutically targetable, as a small molecule inhibitor of NSD2 is being tested in an early-phase clinical trial (NCT05651932) for relapsed t(4,14) translocation-positive multiple myeloma. In combination with our findings, these observations suggest that NSD2 and its relatives play much broader roles in tumor plasticity than previously suspected and represent attractive therapeutic targets.

## Acknowledgements

We are grateful to Steve Henikoff for advice on CUT&Tag analysis. We thank Caisheng Lu in the HICCC Flow Cytometry Core Facility, which is supported in part by the Cancer Center Support Grant P30CA013696. We also thank Erin Bush and Peter Sims for help with single-cell sequencing in the Columbia Genomics and High Throughput Screening Shared Resource of the Herbert Irving Comprehensive Cancer Center. These studies were supported by grants from the NIH P01CA265768 (M.M.S., C.A.-S., C.L.S., M.L., and A.C.), R01CA251527 (M.M.S.), R01CA238005 (M.M.S.), U01CA261822 (M.M.S.), R01CA266040 (K.G.), R01CA253368 (K.G.), R01CA173481 (C.A.-S.), R01CA183929 (C.A.-S.), P50CA211024 (M.L.), R01CA193837 (C.L.S.), P50CA092629 (C.L.S.), U54CA224079 (C.L.S.), R01CA155169 (C.L.S.), P30CA008748 (C.L.S.), R35CA197745 (A.C.), S10OD012351 (A.C.), S10OD021764 (A.C.), R35GM138181 (C.L.), R01DK132251 (C.L.), R01DE031873 (C.L.), by grants from the DOD (PC160357, M.L.), DOD Idea Award (W81XWH20-1-0289, C.L.S), by a DOD Prostate Cancer Research Program Early Investigator Research Award (W81XWH19-1-0337, A.V.), by an Early Career Development Pilot Award through P30CA013696 (A.V.), by the T.J. Martell Foundation (M.M.S.), by the Prostate Cancer Foundation (M.M.S., M.L.), and the Howard Hughes Medical Institute (C.L.S).

## Author contributions

J.J.L. and M.M.S. conceived and designed the overall study, and C.L. designed the epigenetic analyses. J.J.L. established NPPO organoids and performed most of the organoid experiments. A.V. and J.J.L. designed and A.V. performed the computational analyses. M.Z. and C.A.-S. generated mice for NPPO organoid establishment. M.A.R. evaluated mouse organoid histopathology. Z.S. and C.L.S. established and performed analyses of TKO organoid lines. J.J.L., D.K., X.C., and C.L. performed and analyzed epigenetic analyses. B.D.R. and M.L. provided tissue microarrays and J.J.L. F.S., Z.F., T.P., K.G., B.D.R. and M.L. performed staining and analysis of tissue microarrays. J.d.B. analyzed human patient cohorts. Y.C. provided human organoid lines. A.C. supervised computational analyses, and C.L. and M.M.S. supervised experimental analyses. J.J.L. and M.M.S. wrote the initial manuscript, and all authors contributed to its revision.

## Legends for Extended Data Figures

**Extended Data Figure 1.**
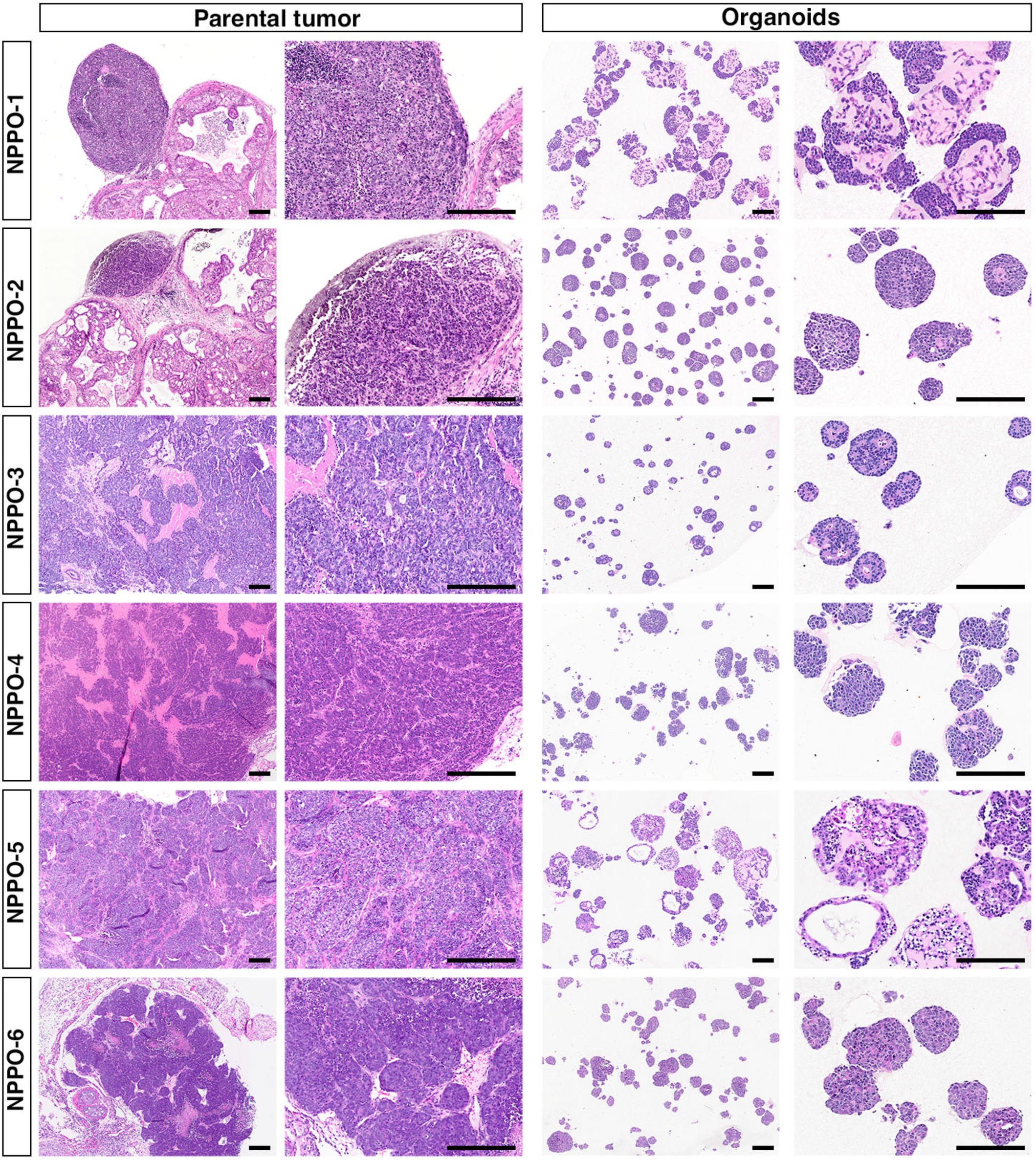
Histology of parental *NPp53* tumors and corresponding NPPO organoid lines. Low- and medium-power views of hematoxylin-and-eosin (H&E) stained sections from the indicated organoid lines at passage 2 and corresponding parental tumors. Scale bars indicate 200 microns.

**Extended Data Figure 2.**
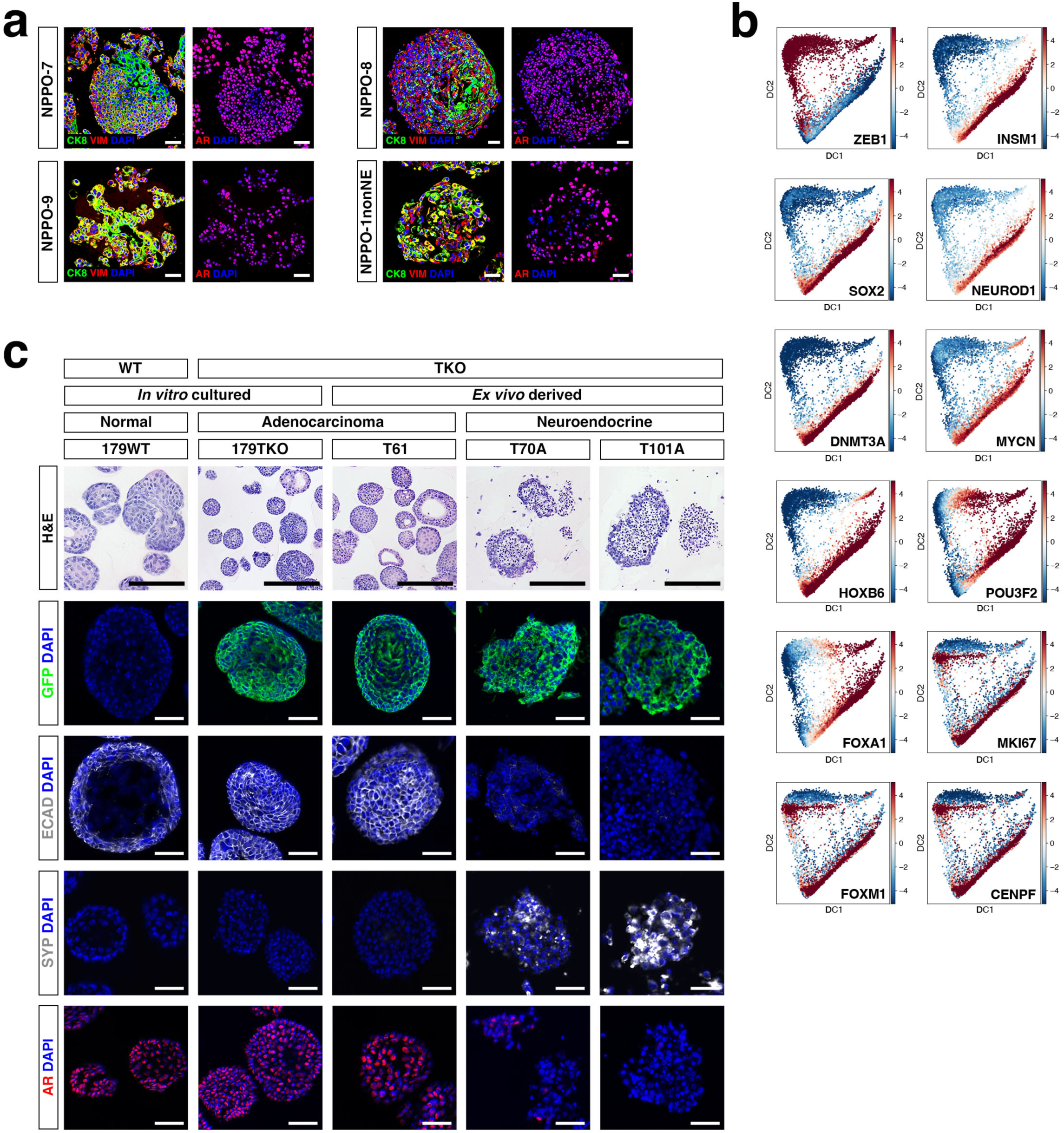
Phenotypes of NPPO and TKO organoid lines. **a,** Immunofluorescence staining for the indicated markers in non-neuroendocrine NPPO organoid lines. AR, Androgen receptor; CK8, Cytokeratin 8; VIM, Vimentin. **b,** VIPER-inferred activity of indicated proteins in the composite neuroendocrine NPPO dataset. **c,** Histological and immunofluorescence analysis of mouse TKO organoids established from *Trp53^flox/flox^*; *Rb1^flox/flox^*; *Pten^flox/flox^* (TKO) mice. WT organoids were prior to infection with Adeno-Cre to induce gene deletion. *Ex vivo* organoids were established from organoid grafts that were subsequently explanted back into organoid culture. GFP expression indicates Cre-mediated recombination. ECAD, E-cadherin. Scale bars in **a,c** indicate 50 microns.

**Extended Data Figure 3.**
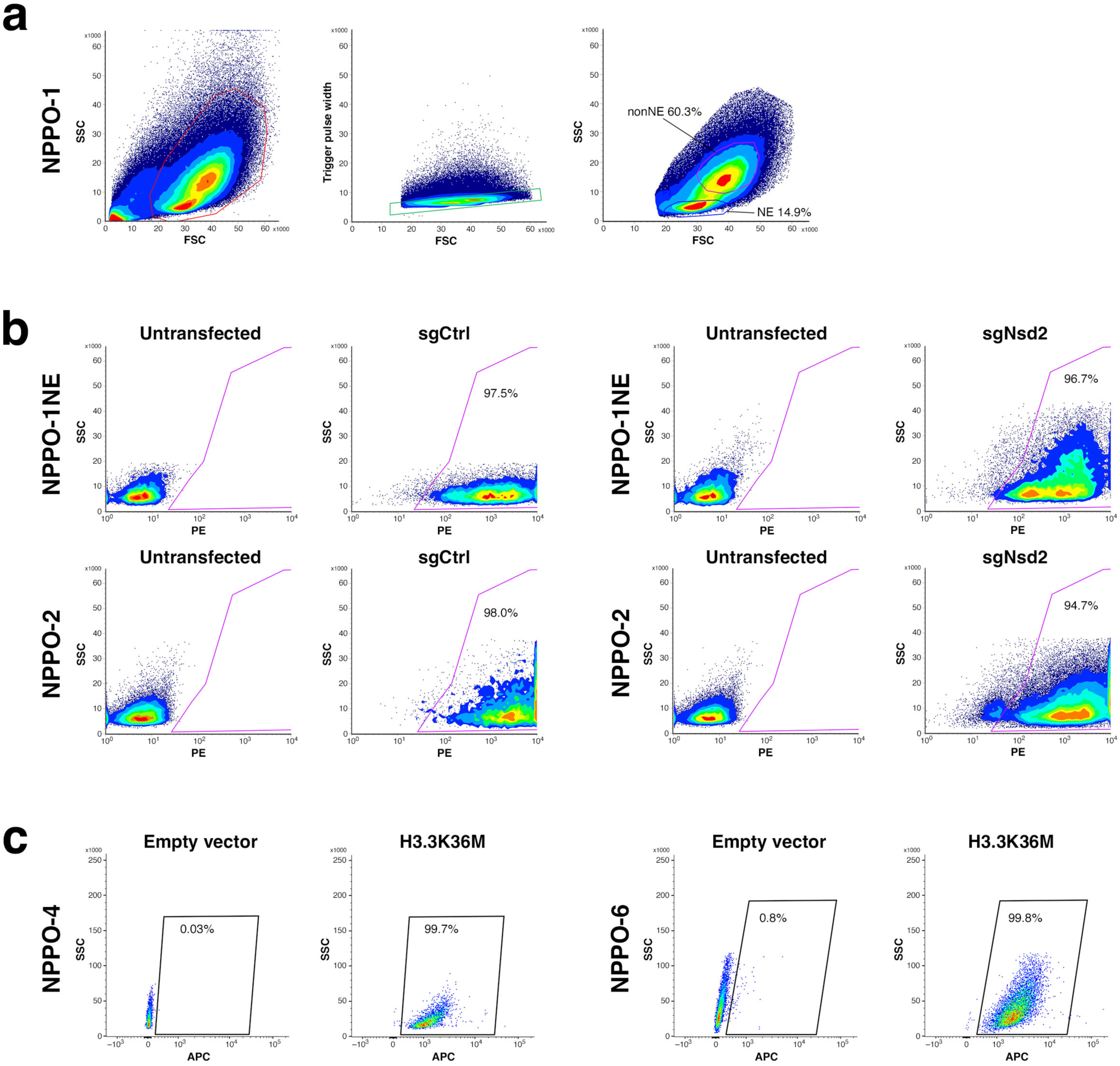
Flow sorting of NPPO organoids. **a,** Sorting strategy for isolation of NPPO-1NE and NPPO-1nonNE sublines from NPPO-1 organoids. **b,** Isolation of sgCtrl and sgNsd2 transfected cells from NPPO-1NE and NPPO-2 organoid lines. **c,** Flow cytometry analysis of H3.3K36M-expressing cells from NPPO-4 and NPPO-6 organoid lines.

**Extended Data Figure 4.**
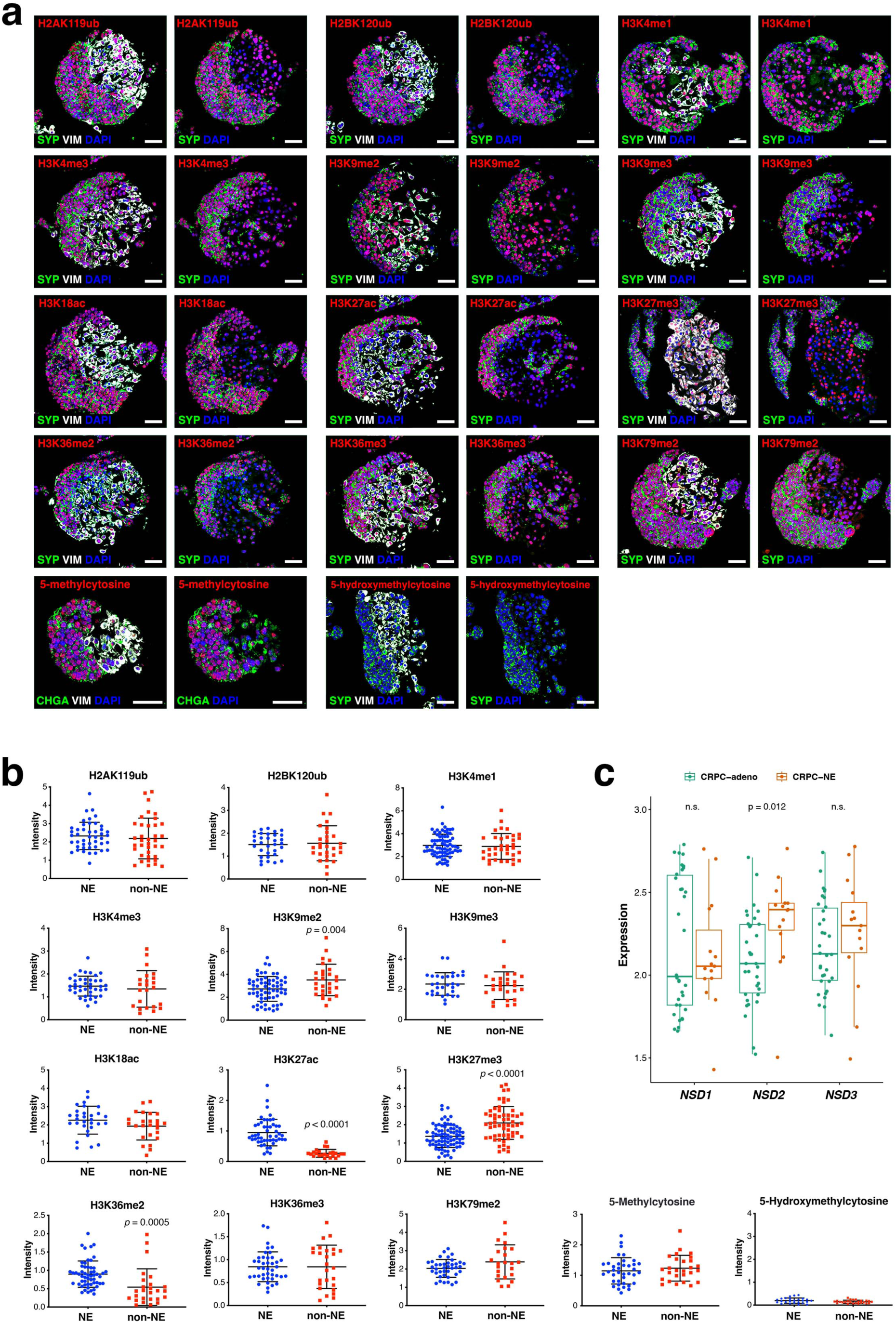
Immunofluorescence screen for differential levels of epigenetic marks. **a,** Immunostaining of indicated epigenetic marks in NPPO-1 organoids. Images are shown in pairs, with and without co-staining for Vimentin (VIM). CHGA, Chromogranin A; SYP, Synaptophysin. Scale bars indicate 50 microns. **b,** Quantitation of epigenetic mark levels, comparing fluorescence intensity in neuroendocrine (NE) and non-neuroendocrine (non-NE) cells in three replicate experiments. Mean fluorescence intensities were compared by unpaired t-tests (two-tailed P value); comparisons lacking p-values were not significant. **c,** Analysis of NSD1, NSD2, and NSD3 expression levels (log_10_ transformed count/million (CPM)) in samples of CRPC-adeno (n=34) and CRPC-NE (n=15) based on data from ^23^. Unpaired t test (two tailed P value) was used for comparison between two groups.

**Extended Data Figure 5.**
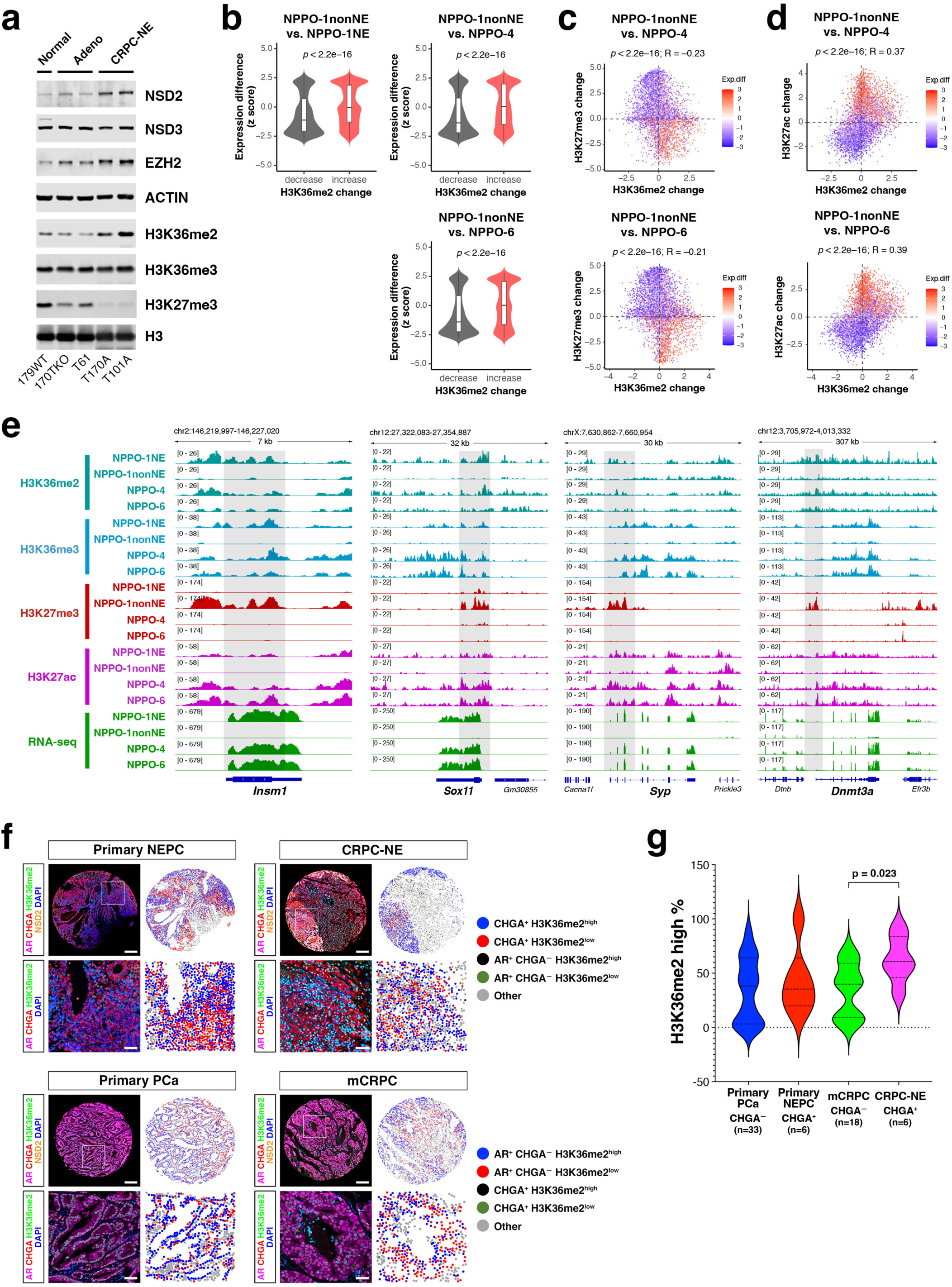
Analysis of histone marks in neuroendocrine organoid lines. **a,** Western blot analysis of indicated proteins and histone marks in TKO organoid lines with normal, adenocarcinoma, and CRPC-NE phenotypes; note that a lane between T61 and T170A was cropped out. **b,** Violin plots showing that genome-wide differential H3K36me2 peaks in NPPO-1nonNE versus NPPO-1NE, NPPO-1nonNE versus NPPO-4, or NPPO-1nonNE versus NPPO-6 are correlated with differential gene expression. Unpaired t test (two-tailed P value) was used to compare differences between two groups. **c,d,** Correlation of genome-wide differential H3K36me2 and H3K27me3 marks (**c**) or H3K36me2 and H3K27ac marks (**d**) with differential gene expression levels in NPPO-1nonNE versus NPPO-4 or NPPO-6. One-way ANOVA was used to determine differences between groups. **e,** Genome browser view of CUT&Tag signals for H3K36me2, H3K36me3, HeK27ac, and H3K27me3 together with bulk RNA-seq reads at *Insm1, Sox11, Syp,* and *Dnmt3a* loci in the indicated organoid lines. **e,f,** Analyses of H3K36me2 levels in a prostate cancer tissue microarray. **f,** Five-color overlay images of representative tissue cores and high-power magnification of four-color images. Spatial plots show the enrichment of H3K36me2^high^ cells among CHGA^+^ neuroendocrine tumor cells or among AR^+^/CHGA^−^ adenocarcinoma. Scale bars indicate 50 microns. **g,** Violin plot for the percentage of H3K36me2^high^ cells in CHGA^+^ neuroendocrine tumor cells or AR^+^/CHGA^−^ tumor cells in each patient. Unpaired t test (two-tailed) was used for comparison between different patient groups.

**Extended Data Figure 6.**
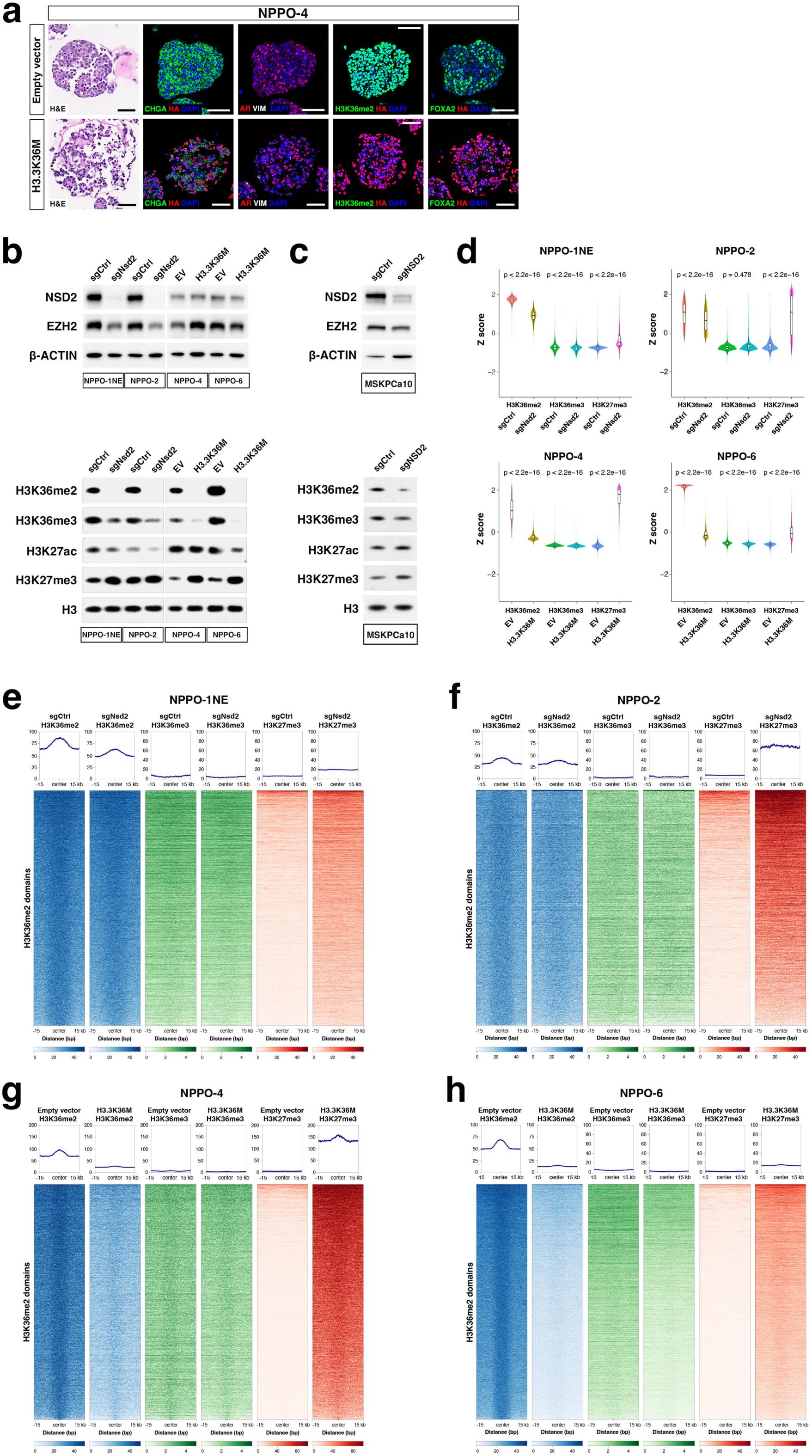
Analysis of histone marks following NSD2 inhibition. **a,** Hematoxylin-and-eosin (H&E) staining and immunofluorescence staining of sections from NPPO-4 organoids after transfection of the oncohistone H3.3K36M or control (EV, empty vector). AR, Androgen receptor; CHGA, chromogranin A; HA, Hemagglutinin tag; VIM, Vimentin. Scale bars indicate 50 microns. **b,** Western blot analysis of NSD2 and EZH2 proteins *(top)* and of H3K36me2, H3K36me3, H3K27ac, and H3K27me3 levels *(bottom)* in control (sgCtrl) and *Nsd2* knock-out (sgNsd2) NPPO-1NE and NPPO-2 organoids, or in control (EV) and H3.3K36M-transfected NPPO-4 and NPPO-6 organoids. **c,** Western blot analysis for control (sgCtrl) and *NSD2* knock-out (sgNSD2) MSKPCa10 organoids. **d,** Violin plots showing quantitative comparison of H3K36me2, H3K36me3 and H3K27me3 CUT&Tag signals at genomic regions marked by H3K36me2 between sgCtrl and sgNsd2 or between EV and H3.3K36M-transfected NPPO organoids. Wilcoxon rank-sum test was used to compare two independent samples. **e-h,** Heatmaps of CUT&Tag signals for the indicated histone marks at genomic regions marked by H3K36me2, shown for NPPO-1NE (**e**), NPPO-2 (**f**), NPPO-4 (**g**), and NPPO-6 (**h**).

**Extended Data Figure 7.**
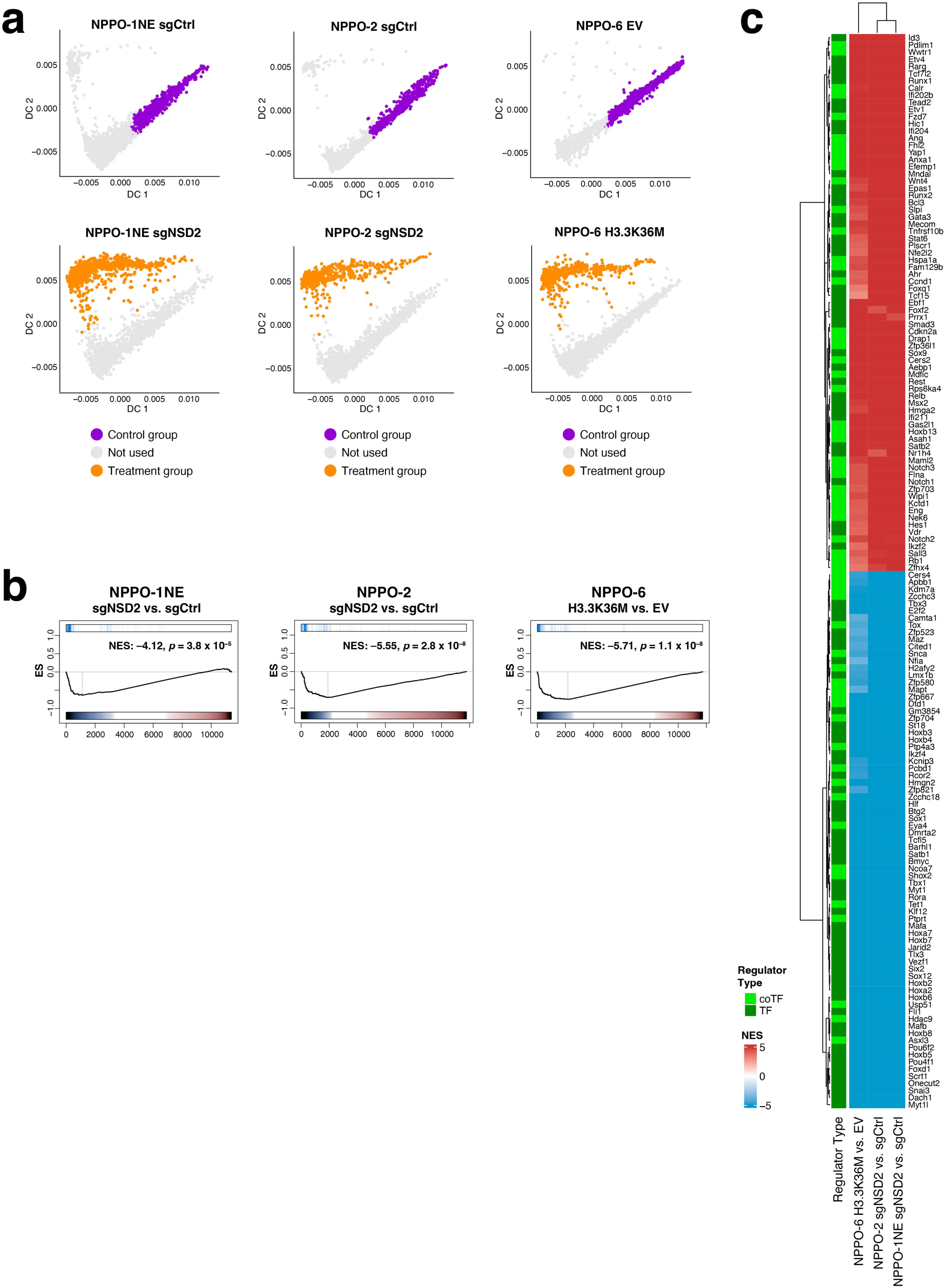
Differential activity of genes following NSD2 inhibition. **a,** Selection of cell populations for comparison from Cluster 3 *(top)* or Cluster 1 *(bottom)* from NPPO-1NE and NPPO-2 sgCtrl and sgNsd2 organoids or from NPPO-6 EV and H3.3K36M organoids. **b,** Single-cell Gene Set Enrichment Analysis (scGSEA) plots of gene expression signatures computed as differential between cell clusters indicated in panel **a**, showing enrichment of genes with decreased H3K36me2 levels identified by CUT&Tag analysis after *Nsd2* targeting or H3.3K36M expression. **c,** Heat map shows transcription factors (TFs) and co-transcription factors (co-TFs) prioritized by VIPER that are differentially active after NSD2 inhibition or H3.3K36M expression.

**Extended Data Figure 8.**
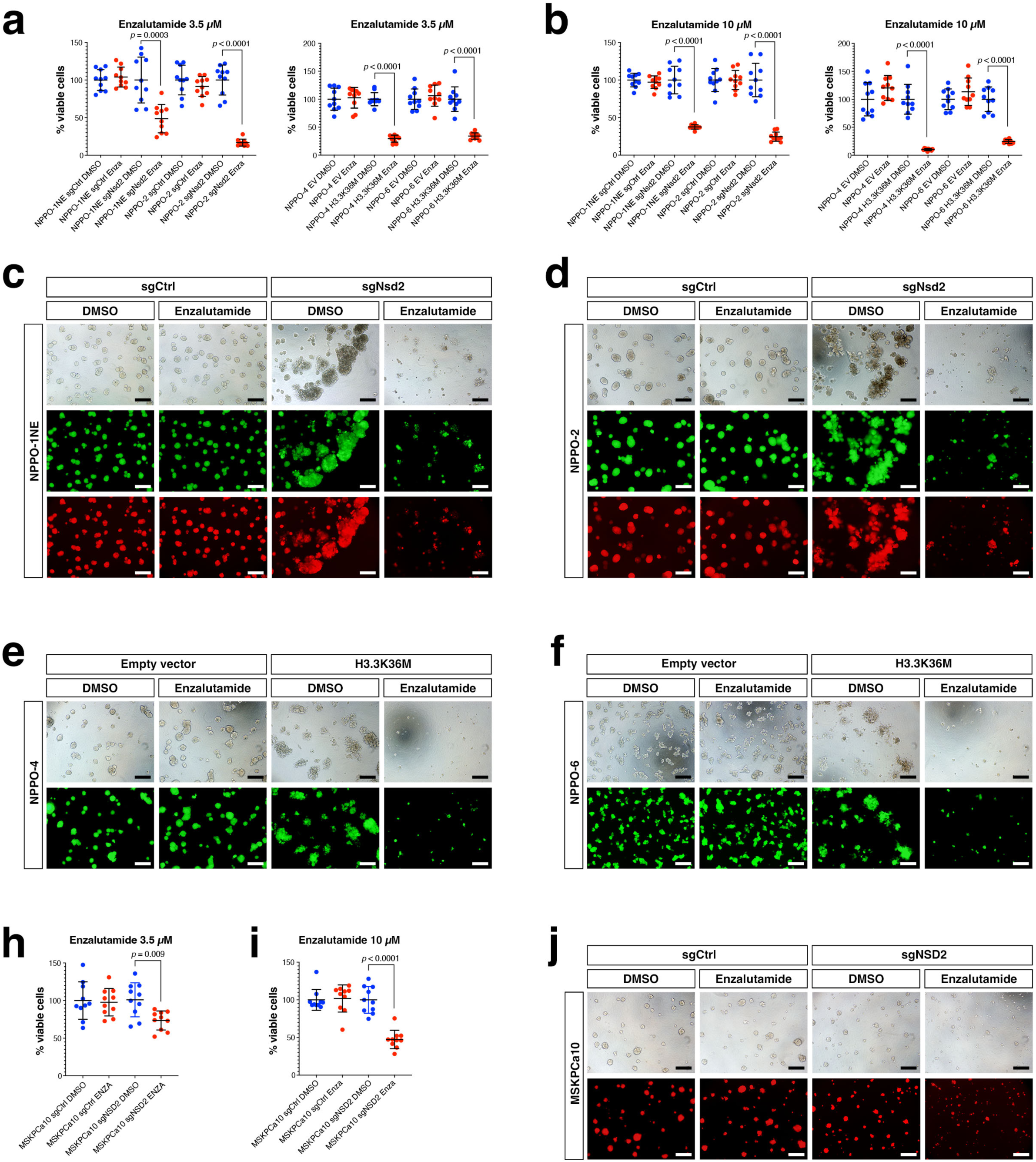
Organoid response to combined NSD2 inhibition and enzalutamide treatment. **a,b,** Growth of *Nsd2* knock-out (sgNsd2) or control (sgCtrl) NPPO-1NE and NPPO-2 organoids (**a**), control (EV, empty vector) or H3.3K36M-transfected NPPO-4 and NPPO-6 organoids (**b**), Experimental values were normalized to DMSO controls and shown as percentage of viable cells (n=10 biological replicates). p-values were calculated using unpaired t-tests (two-tailed). **c-f,** Whole-mount images of organoids following *Nsd2* knock-out or H3.3K36M expression treated with either DMSO control or 10 µM enzalutamide. All NPPO organoid lines express YFP (green) due to the Cre reporter in the *NPp53* mouse model; the NPPO-1NE, NPPO-2 organoids additionally express RFP following sgCtrl or sgNSD2 transfection. **h,i,** Growth of sgNSD2 or sgCtrl MSKPCa10 organoids treated with 3.5 µM (**h**) or 10 µM enzalutamide (**i**) or DMSO control. **j,** Whole-mount images of MSKPCa10 organoids following *NSD2* knock-out or H3.3K36M expression treated with either DMSO control or 10 µM enzalutamide. These organoids express RFP following sgCtrl or sgNSD2 transfection.

**Extended Data Figure 9.**
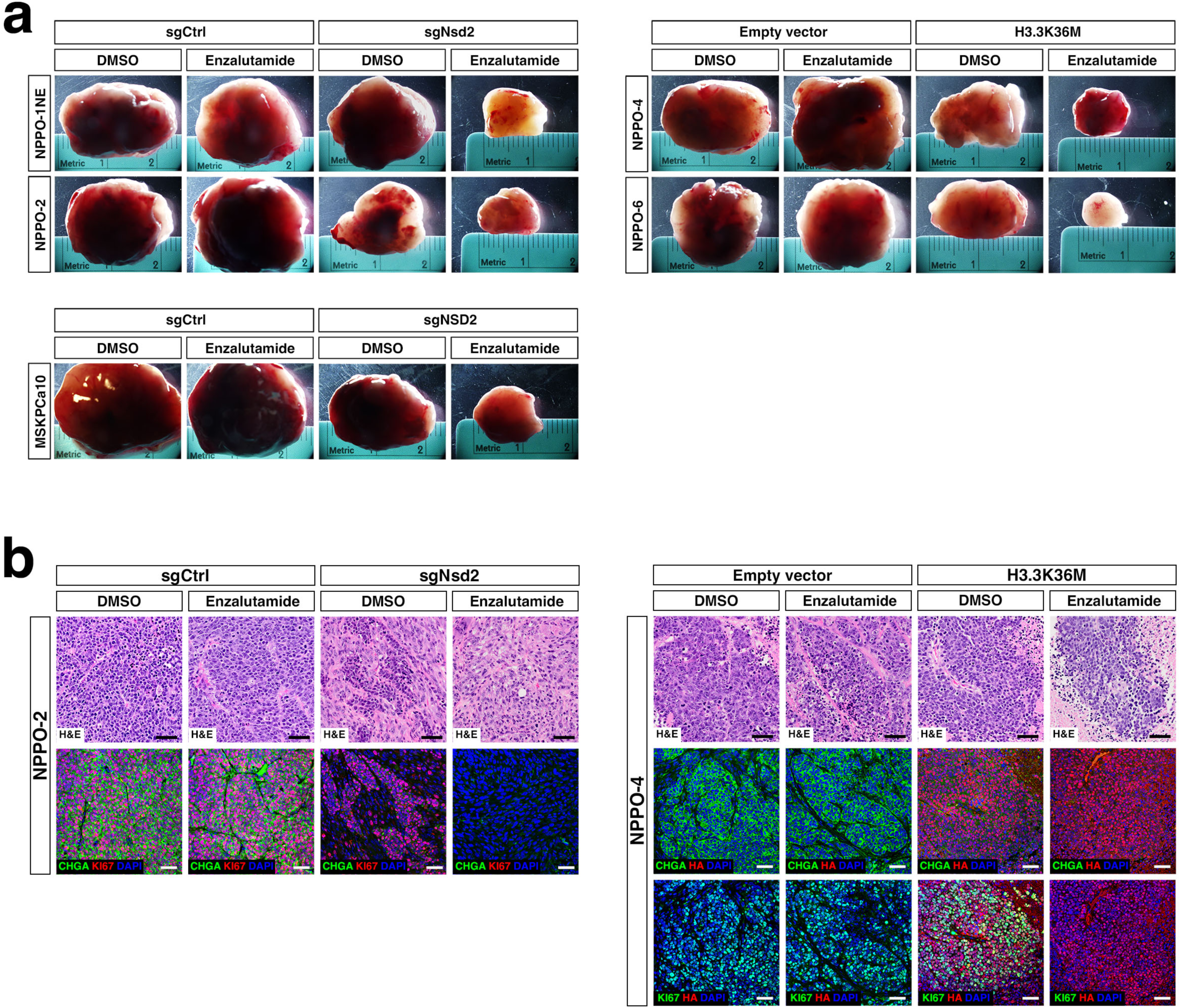
Graft response to combined NSD2 inhibition and enzalutamide treatment. **a,** Whole-mount images of grafts following subcutaneous implantation of NPPO-1NE, NPPO-2, and MSKPCa10 control or *NSD2* knock-out organoids *(left)* or NPPO-4 and NPPO-6 control or H3.3K36M-transfected organoids *(right)* into NOD/SCID immunodeficient mice treated with DMSO control or enzalutamide. **b,** H&E and immunofluorescence analysis of sections from NPPO-2 *(left)* and NPPO-4 *(right)* grafts. CHGA, Chromogranin A; HA, Hemagglutinin tag. Scale bars in **b** indicate 50 microns.

## Supplementary Tables

**Supplementary Table 1.** Mice used for establishment of organoid lines.

**Supplementary Table 2.** Histopathology of mouse and human organoids and grafts.

**Supplementary Table 3.** Single-cell RNA-seq, ATAC-seq, and multiome datasets analyzed in this study.

**Supplementary Table 4.** Samples included on human prostate cancer tissue microarray.

**Supplementary Table 5.** Antibodies used in this study.

**Supplementary Table 6.** Primers used for CUT&Tag library construction.

## Methods

### Mouse procedures

The *Nkx3.1^CreERT2/+^; Pten^flox/flox^; TrpP53^flox/flox^; Rosa26-EYFP (NPp53)* mice were maintained on a mixed C57BL/6-129Sv background and have been previously described^7^. Tamoxifen induction was performed in mice at 3-5 months of age by oral delivery of tamoxifen (Sigma; 100 mg/kg/day in corn oil) for 4 consecutive days as described previously^69^. The survival time of tumor-bearing mice in this study ranged from 228 to 435 days after tamoxifen induction (Supplemental Table 1). All procedures followed protocols approved by the Institutional Animal Care and Use Committee (IACUC) at Columbia University Medical Center.

### Establishment and maintenance of mouse prostate organoids

Tumor tissues from *NPp53* mice were cut into two pieces, with half fixed in 10% formalin for paraffin embedding, and the other half used for organoid establishment. Tissues were minced with scissors in 0.2% collagenase IV (Thermo Fisher Scientific 17104019) and incubated at 37°C for 30 min, followed by neutralization with 1:10 Hank’s buffer (STEMCELL Technologies 37150) supplemented with 10 µM Y-27632 (STEMCELL Technologies) and 5% charcoal-stripped fetal bovine serum (CS-FBS; Gemini 100-119). After centrifugation at 1000 rpm for 10 min, pellets were incubated with prewarmed TrypLE (Thermo Fisher Scientific 12605010) at 37°C for 10 min. The cell suspension was then neutralized 1:10 with PBS, passed through a 100 µM cell strainer (Corning 352360), and spun down at 1000 rpm for 10 min. Prior to plating, cell numbers were counted in a TC20 automated cell counter (Bio-Rad).

Cells were resuspended in organoid culture media supplemented with 10 µM Y-27632, 10 µM A83-01 (Tocris 2939), and 5% Matrigel (Corning 354234) and plated at a seeding density of approximately 50,000 cells/well in 96-well ultra-low attachment microplates (Corning 3474). Organoid culture medium consisted of hepatocyte culture medium (Corning 355056), 5% CS-FBS, 1X GlutaMAX™ supplement (Thermo Fisher Scientific 35050061), 5 ng/ml EGF, 100 µg/ml primocin (Invivogen ant-pm-1) and 100 nM dihydrotestosterone (DHT), as previously described^70^. For heterogeneous NE organoids, such as NPPO-1 and NPPO-5, organoid culture medium was replaced every 4 days. For homogeneous NE organoids, such as NPPO-1NE, NPPO-2, NPPO-4 and NPPO-6, we used NE organoid culture medium, which was identical to the organoid culture medium except that no EGF was added; the medium was replaced every 4 days.

For passaging, organoids were collected by centrifugation at 1000 rpm for 1 min, followed by addition of 1 ml pre-warmed TrypLE for 10 min at 37°C for cell dissociation. After neutralization with 10 ml PBS, cells were spun down and counted, with approximately 50,000 cells plated per well in 96-well ultra-low attachment microplates (Corning 3474). To generate cryopreserved stocks, organoids were frozen in 90% CS-FBS and 10% DMSO and stored in liquid nitrogen. We considered NE organoid lines to be successfully established when they could be stably passaged, cryopreserved, and recovered without loss of NE phenotypes.

Complete details on the isolation and characterization of the TKO organoids (Extended Data Figures 2C and 5A) will be reported separately.

### Human prostate tumor organoids

MSKPCa10 organoids have been previously described^52^ and were generously provided by Dr. Yu Chen (Memorial Sloan Kettering Cancer Center). Human prostate tumor organoids were maintained in 60% Matrigel and 40% human NE culture medium, which was replaced every other day. Human NE culture medium consisted of hepatocyte culture medium, 5% CS-FBS, 1X GlutaMAX™, 5 ng/ml EGF, 100 µg/ml primocin and 10 nM DHT.

### Hematoxylin-eosin staining

For tissue processing and embedding, organoids were fixed in 10% formalin (Thermo Fisher Scientific, SF100-4) for 1 h, washed once with PBS, placed in rat tail collagen I (Corning 354249) and incubated at 37°C for 30 min. The collagen button was then put into a biopsy cassette (Thermo Fisher Scientific 15182705E) and fixed in 10% formalin for 24 h. After replacing the formalin with 70% ethanol, the cassettes were put into an automated tissue processor for tissue processing and embedding.

Paraffin-embedded blocks were sectioned into 5 µm sections using microtome and dried by onto microscope slides at RT. Paraffin sections were baked at 65°C for 15 minutes prior to deparaffinization with three changes of xylene, 5 min each. The slides were hydrated through 100%, 95%, and 95% ethanol, 5 min each, rinsed in tap water for 2 min, and incubated in Gill™ Hematoxylin 3 (Epredia 72611) for 3-30 min. Slides were rinsed in tap water and dipped 3-5 times in 0.5% Acid Alcohol (Leica Biosystems 3803651), followed by rinsing in tap water and bluing with Scott’s Tap Water (Electron Microscopy Sciences 2607007) for 5 min. After rinsing in tap water, slides were incubated in 95% ethanol for 5 min and counterstained with Eosin (Poly Scientific S1761GL) for 1-3 min. Slides were passed through 70%, 95%, and 100% (3x) ethanol, rinsed 3 times in xylene, and coverslipped with mounting medium (StatLab, MMC0126). Images were captured using a Olympus BX 61 VS Slide Scanner.

### Immunofluorescence staining

Paraffin sections (5 µm) were dried onto microscope slides at RT, incubated at 65°C for 15 min prior to deparaffinization through three changes of xylene (5 min each), hydrated in 100%, 95%, 95%, 75% ethanol (5 min each), and washed in tap water for 2 min. Antigen retrieval was performed by immersion in boiling citrate buffer (pH 6) for 10 min, cooling to RT for 30 min, and incubation in Milli-Q water at RT for 10 min. Sections were permeabilized with 0.5% Triton X-100 in PBS (MilliporeSigma 11332481001) for 10 min and blocked in 10% goat serum for 1 hr. Diluted primary antibodies (Supplemental Table 5) were added to sections and incubated overnight at 4°C. The next day, sections were washed with PBS three times, 15 min each, and incubated with secondary antibodies at RT for 1 hr. After washing with PBS three times, 15 min each, nuclei were stained with DAPI (Thermo Fisher Scientific D1306) for 5 min. Slides were washed with PBS and mounted with VECTASHIELD® Antifade Mounting Medium (Vector Laboratories H-1200-10). Images were captured using a Leica TCS SP5 Confocal Laser Scanning Microscope (Leica Microsystems) using Leica Application Suite Advanced Fluorescence (LAS AF).

### Single-cell RNA sequencing

For scRNA-seq, NPPO-1, NPPO-2, NPPO-4, NPPO-5, and NPPO-6 organoids were analyzed at passage 2. Organoids were dissociated by incubation with pre-warmed TrypLE at 37°C for 10 min, neutralization with 1:10 5% CS-FBS/PBS, centrifugation at 1000 rpm for 1 min, and filtration three times through a 40 µM cell strainer (Corning 431750). Cells were spun at 1000 rpm for 5 min, pellets were dissociated with 5% CS-FBS/PBS, and resuspended at 1000 cells/µl after counting using a Countess II FL Automated Cell Counter. Libraries were prepared using a Chromium Next GEM Single Cell 3ʹ Reagent Kit v3.1 by the Columbia University Single Cell Analysis Core. Approximately 5,000 cells were loaded onto the Chromium Controller (10X Genomics) to generate Gel Beads-in-emulsion (GEMs), and barcoded, full-length cDNA from poly-adenylated mRNA was generated and amplified via PCR. Chromium Gene Expression libraries were prepared for paired-end sequencing, and scRNA-seq data were processed with Cell Ranger software (10x Genomics, version 2.1) by the Columbia University Single Cell Analysis Core.

### Single-nucleus ATAC sequencing

NPPO-1 organoids at passage 8 were dissociated using TrypLE, passed through a 40 µM cell strainer three times, and cell numbers quantified using an automated cell counter. Approximately 1 x 10^6^ cells in 0.04% BSA/PBS (Miltenyi Biotec 130-091-376) were used for single nuclei isolation. Cells were spun at 1000 rpm for 5 min at 4°C, dissociated for 5 min with 100 µl ice-cold lysis buffer, neutralized with 1 ml chilled wash buffer, spun at 1000 rpm for 5 min at 4°C, and resuspended in chilled nuclei buffer included in the Chromium Single Cell ATAC Library Kit (10X Genomics). Wash buffer contained 10 mM Tris-HCl (pH 7.4), 10 mM NaCl, 3 mM MgCl_2_, 1% BAS, 0.1% Tween-20 in nuclease-free water. Lysis buffer contained 10 mM Tris-HCl (pH 7.4), 10 mM NaCl, 3 mM MgCl_2_, 0.1% Tween-20, 0.1% Nonidet P40 Substitute, 0.01% Digitonin, 1% BAS in nuclease-free water. Nuclei concentration was determined using a Countess II FL Automated Cell Counter. Approximately 5,000 nuclei were loaded onto the Chromium Controller at the Columbia University Single Cell Analysis Core. snATAC-seq libraries were prepared following the manufacturer’s instructions (Chromium Next GEM Single Cell ATAC Reagent Kits v1.1, 10X Genomics). Briefly, nuclei were transposed and partitioned into Gel Beads-in-emulsion (GEMs). 10x Barcodes were added to index the transposed DNA of each individual nucleus. Libraries were generated via PCR and sequenced on the Illumina NextSeq 550 platform. Paired-end sequencing data were processed with Cell Ranger software (10x Genomics, version 2.1) by the Columbia University Single Cell Analysis Core.

### Isolation of NE and non-NE cells from NPPO-1 organoids

To sort NE and non-NE populations, NPPO-1 organoids at passage 2 were incubated with prewarmed TrypLE at 37°C for 10 min, neutralized with 1:10 PBS and 5% CS-FBS, spun at 1000 rpm for 1 min, resuspended with PBS, and dissociated into single cells by gentle pipetting. The cells were filtered three times through a 40 µm cell strainer (Corning 431750), spun at 1000 rpm for 5 min, and resuspended with PBS and 2% CS-FBS. After filtering through a Falcon tube with 35 µm strainer cap (Corning 352235), cell numbers were counted in a TC20 automated cell counter, and the volume adjusted to a final cell concentration of 5,000 cells/µl.

Flow sorting was performed on an BD Influx™ cell sorter (BD Biosciences) in the Flow Cytometry Core of the Columbia Center for Translational Immunology (CCTI). Gating by forward scatter (FSC) and side scatter (SSC) was used to exclude debris, and doublets were excluded by gating on trigger pulse width against FSC height. Individual NE and non-NE tumor cells were sorted based on scatter parameters. as NE tumor cells have less internal complexity (granularity) than non-NE tumor cells and exhibit lower intensity SSC. Flow sorting data were collected and analyzed using BD FACS™ Software (BD Biosciences, version 1.2.0.142). Cell purity was assessed following flow sorting by single-cell RNA sequencing.

### Lineage-tracing in organoids

Flow-sorted NE cells from NPPO-1 organoids were maintained in NE organoid culture medium with 5% Matrigel. Half of the flow-sorted non-NE cells were used for scRNA-seq (Columbia University Single Cell Analysis Core), immediately after sorting. The other non-NE cells were transfected with H2B-RFP (Addgene 26001) lentivirus. Approximately 70% of non-NE cells were labeled with H2B-RFP at 3 days after transfection, in the absence of antibiotic selection. On day 7, the cells were digested with TrypLE and passed three times through a 40 µm cell strainer to ensure a single cell suspension, following by cell counting. For co-culture, H2B-RFP labeled non-NE cells were seeded together with NE cells at a ratio of 2:3 in 96-well ultra-low attachment microplates; as a control, H2B-RFP labeled non-NE cells were seeded alone. The resulting organoids were cultured in NE organoid culture medium with 5% Matrigel, and analyzed at passage 4 by immunostaining and scRNA-seq.

### Imaging of histone and DNA modifications

To screen for differential expression of histone modifications between NE and non-NE tumor cells in NPPO-1 organoids, we performed immunofluorescence staining with antibodies detecting the NE markers Synaptophysin (SYP) or Chromogranin A (CHGA), the non-NE marker Vimentin (VIM), and various histone/DNA modifications (Supplemental Table 5). Images were captured using a Leica TCS SP5 confocal laser scanning microscope and images acquired with Leica Application Suite Advanced Fluorescence (LAS AF). Fluorescence intensity was measured using ImageJ (NIH; version 1.52K) using three parameters: area, integrated density (IntDen), and mean gray value. Background was measured from a region that had no fluorescence on the same image. Intensity was calculated using the formula: Intensity = IntDen – (Area × Mean fluorescence of background readings) using measurements collected from three independent organoids, and graphed using Prism 9 (GraphPad software version 9.3.1). An unpaired t-test was used to compare means, and *P* values calculated from a two-tailed t test.

### Histone extraction and Western blotting

Histone extract lysates were prepared by acid-extraction as described previously^71^. Briefly, cells were resuspended in hypotonic lysis buffer (10 mM Tris–Cl pH 8.0, 1 mM KCl, 1.5 mM MgCl_2_ and protease inhibitor cocktail (MilliporeSigma 11697498001)), and incubated on a rotator for 30 min at 4°C. Nuclei were isolated by centrifugation (16,000 g for 10 min at 4°C) and resuspended in H_2_SO_4_. After overnight incubation on rotator at 4°C, debris was removed by centrifugation (16,000 g for 10 min at 4°C) and histones precipitated from the supernatant using TCA (MilliporeSigma T6399). Purified histones were washed with cold acetone and resuspended in H_2_O. Samples were quantified using Bradford assay (Bio-Rad 5000205), and protein lysates were prepared using SDS lysis buffer (Thermo Fisher Scientific LC2676). Whole cell lysates were prepared by resuspending cells in SDS lysis buffer, sonication for 10 seconds twice, and boiling for 8 min. Protein lysates were resolved on 3-8% (Thermo Fisher Scientific EA0375BOX) or 4-12% gradient SDS-PAGE gels (Thermo Fisher Scientific NP0321BOX), transferred to nitrocellulose membrane, blocked in 5% non-fat milk in PBS plus 0.5% Tween-20, probed with primary antibodies (Supplemental Table 5), and detected with horseradish peroxidase (HRP)-linked anti-rabbit IgG (Cell Signaling Technology 7074) or anti-mouse IgG (Cell Signaling Technology 7076). Blots were imaged using a ChemiDoc MP Imaging system (Bio-Rad 17001402) or exposed to X-ray film (Research Products International 248300).

### Bulk RNA-sequencing

Organoids were dissociated into single cells using TrypLE, and total RNA isolated using MagMAX™-96 for Microarrays Total RNA Isolation Kit (Thermo Fisher Scientific AM1839). Briefly, cells were lysed in 1 ml TRI reagent per 5 million cells, and incubated for 5 min at RT. The homogenate was mixed with 0.1 volume 1-bromo-3-chloropropane (MilliporeSigma B9673), incubated at RT for 5 min, and centrifuged at 12,000 g for 10 min at 4°C. 100 μl of the aqueous phase was transferred to an 8-strip EpiCypher tube (DNase/RNase-free), and 50 μl of 100% isopropanol (MilliporeSigma I9516) added with shaking for 1 min. 10 μl of RNA Binding Beads were added, followed by shaking for 3 minutes. The RNA Binding Beads were then captured on a magnetic stand and the beads washed twice with 150 μl Wash Solution. The beads were dried for 2 min, and RNA eluted in 50 μl of Elution Buffer. All samples were DNase-treated prior to library construction. Quality assessment was performed using the 4200 TapeStation system (Agilent), with a median RNA integrity number of 9.2 (range 8.7 to 9.8). Libraries were generated using the Illumina Stranded mRNA Prep, and 150-bp paired-end sequencing was performed to a minimum of 20 million reads per sample on the Illumina HiSeq platform.

### Cleavage Under Targets and Tagmentation (CUT&Tag)

Prior to cell preparation, 10 µl/sample concanavalin A (ConA)-coated magnetic beads (Bangs Laboratories BP531) were transferred into a 1.5 ml LoBind tube (Eppendorf 22431021) and mixed by gentle vortexing with 0.4 ml binding buffer (20 mM HEPES (pH 7.5; Thermo Fisher Scientific 15630080), 10 mM KCl (MilliporeSigma 60142), 1 mM CaCl_2_ (MilliporeSigma 21115) and 1 mM MnCl_2_ (MilliporeSigma M1787) in nuclease-free water). The tubes were placed on a magnetic stand (Thermo Fisher Scientific 12321D) to clear and remove liquid, with the wash repeated once with binding buffer.

For experiments requiring normalization of input DNA, 10,000 Drosophila Schneider 2 (S2) cells (Thermo Fisher Scientific R69007) were combined with 200,000 experimental cells at a ratio of 1:20. S2 cells were maintained in Schneider’s Drosophila Medium (Thermo Fisher Scientific 21720024), 10% heat-inactivated FBS (Gemini 100-106) and 100 µg/ml primocin. S2 cells were plated in 96-well ultra-low attachment microplates at a seeding density of approximately 100,000 cells/well and maintained at 28°C without CO_2_.

Organoids were dissociated into single cells using TrypLE, counted using a TC20 automated cell counter, and 200,000 cells/sample used for profiling. After centrifugation at 1000 rpm at RT, cell pellets were washed once with 1 ml wash buffer (20 mM HEPES (pH 7.5), 150 mM NaCl, 0.5 mM spermidine (MilliporeSigma S2501), and cOmplete™, EDTA-free protease inhibitor cocktail (MilliporeSigma 11873580001) in nuclease-free water). Cell pellets were resuspended with 10 µl ConA beads in 0.4 ml wash buffer and incubated for 10 minutes at RT. The tubes were then placed on a magnetic stand (New England Biolabs S1515S) to clear and remove the liquid, with pellets resuspended with 50 µl ice-cold Dig-wash buffer (wash buffer with 0.05% digitonin), and placed on ice for 10 min to lyse the cells. The samples were then transferred to an 8-strip tube (EpiCypher 10-0009). Each sample was incubated with 0.5 µg primary or IgG control antibody (Supplemental Table 5) and placed on a nutator at 4°C overnight. The next day, samples were incubated with 1 µl secondary antibody in 50 µl ice-cold Dig-wash buffer on a nutator at RT for 1 hr. After washing with Dig-Wash buffer three times, beads were resuspended with 50 µl Dig-300 buffer (20 mM HEPES (pH 7.5), 300 mM NaCl, 0.5 mM Spermidine, 0.01% Digitonin, cOmplete™, EDTA-free Protease Inhibitor Cocktail in nuclease-free water) containing 2.5 µl pA-Tn5 adapter complex (EpiCypher 15-1017) and placed on a nutator at RT for 1 hour. After washing three times with Dig-300 buffer, the beads were resuspended in 150 µl Tagmentation buffer (Dig-300 buffer with 10 mM MgCl_2_) and incubated at 37°C for 1 hr.

To stop tagmentation and solubilize DNA fragments, 5 µL 0.5 M EDTA (Thermo Fisher Scientific AM9260G), 1.5 µl 10% SDS (MilliporeSigma 71736), 1.25 µl 20 mg/ml Proteinase K (Thermo Fisher Scientific EO0491) was added to each sample and incubated at 37°C overnight. The next day, phenol:chloroform:isoamyl alcohol (25:24:1, v/v; Thermo Fisher Scientific 15593049) was used to extract nucleic acids. DNA was precipitated with 100% ethanol (MilliporeSigma E7023) and dissolved in 25 μl RNase-free water. To remove RNA contamination prior to library preparation, DNA samples were incubated with 25 µg/ml RNase A (Thermo Fisher Scientific EN0531) at 37°C for 10 min. Library DNAs were amplified by PCR using Illumina universal i5 primers, Nextera barcoded i7 primers (Supplemental Table 6), and NEBNext HiFi 2x PCR Master mix (New England Biolabs M0541). The PCR conditions were: cycle 1: 72°C for 5 min; cycle 2: 98°C for 30 sec; cycle 3: 98°C for 10 sec; cycle 4: 63°C for 10 sec, repeating 14 times, followed by 72°C for 1 min and hold at 8°C. Post-PCR clean-up was performed using 1:1 volume Ampure XP beads (Beckman Coulter A63880). Samples were dissolved in 10 mM Tris-HCl (pH 8; MilliporeSigma T2694) for sequencing. Size distribution and concentration of libraries was determined by capillary electrophoresis using the 4200 TapeStation (Agilent). Paired-end sequencing (2×150 bp) was performed on pooled-libraries, with 10-12 million reads per library, using the Illumina HiSeq system.

### CUT&Tag data analysis

CUT&Tag reads were aligned to the mouse reference genome (mm10) and Drosophila (if Drosophila S2 cells were used as spike-in control) using BOWTIE2 (v2.4.2) with options: --very-sensitive-local --no-unal --no-mixed --no-discordant --phred33 -I 10 -X 700. Scaling factors for spike-in normalization was determined by the ratio between the number of reads aligned to the mouse genome and Drosophila genome. Potential PCR duplicates were removed by the MarkDuplicates function of Picard (v2.23.1). Peak calling was performed using SICER2 with IgG input as control. The read counts of peaks across gene loci were counted by featureCounts (v1.6.1). Genomic enrichment of CUT&Tag signal for each histone modification was analyzed by deeptools (v2.0) and visualized using IGV (v2.13.0). RNA-seq reads were aligned to the mouse reference genome (mm10) using HISAT2 (v2.1.0). The mapped reads count of each gene was measured by featureCounts (v1.6.1). The RNA-seq reads count matrix was combined with the CUT&Tag signal reads count matrix for all gene loci in R (v4.1.2).

### Lentivirus production and transfection

Lentiviruses were generated by transfection of 293T cells with the indicated expression plasmid and the psPAX2 (Addgene 12260) and pVSVG (Addgene 14888) packaging vectors at a ratio of 4:2:3, respectively. Viral supernatants were collected at 48, 72, and 96 h after transfection, filtered, and concentrated using Lenti-X Concentrator (Takara Bio 631232). For CRISPR/Cas9 gene knockout, we used the lentiCas9-blast plasmid (Addgene 52962) and a custom vector for sgRNA (U6-sgRNA-EFS-Puro-P2A-TurboRFP in a pLL3-based lentiviral backbone; gift from Scott Lowe). For sgRNA design, the CRISPick platform (BROAD institute) was used. HA-tagged H3.3K36M was overexpressed in the pCDH vector (gift from David Allis). The sgRNAs used in the experiment are: sgControl, 5’ GAG ATA AGC ATT ATA ATT CCT 3’; sgNsd2 (mouse): 5’ TCA GGG TCT CAC AAT TGG GC 3’; sgNSD2 (human): 5’ GCA CCA GCT CAC GTT GAC GT 3’.

For transfection, organoids were incubated with high-titer lentivirus in culture medium supplemented with 8 ug/ml polybrene (MilliporeSigma TR-1003). Medium containing virus was removed on the next day and switched to normal organoid medium with Matrigel. Selection with appropriate antibiotics was performed at 3 days after transfection for 7-14 days.

### Multiplexed staining of tissue microarrays

Primary antibodies were tested on prostate tumor samples to verify the expected pattern of staining and were titrated at 4 concentrations to determine the best signal-to-noise ratio. Multiplexed staining was performed using the Opal 6-plex detection kit (Akoya Biosciences NEL871001KT) on a Bond Rx Research Stainer (Leica Biosystems), adding DAPI as a nuclear marker. Slides were imaged in the Vectra Polaris Automated Quantitative Pathology Multispectral Imaging System (Akoya Biosciences). Exposure times were optimized under the constraint that no pixel saturate the detector.

### Analysis of patient survival curves

To evaluate the association of NSD2 with overall survival in mCRPC, two independent mCRPC biopsy RNAseq cohorts were used: 1) A cohort of 159 mCRPC transcriptomes generated by the SU2C/PCF Prostate Cancer Dream Team; 141 mCRPC transcriptomes from this dataset were used for the survival analyses as survival data was not available from 18 patients. 2) A cohort of 95 mCRPC transcriptomes from patients treated at the Royal Marsden were analyzed; 94 mCRPC transcriptomes were used for the survival analyses as survival data was not available from one patient. Transcriptomes were aligned to the human reference genome (GRCh37/hg19) using TopHat2 (v2.0.7). Gene expression as fragments per kilobase of transcript per million mapped reads (FPKM) was calculated using Cufflinks (v2.2.1). Kaplan-Meier studies evaluated overall survival outcomes.

### Computational analysis of multiplex images

All TMA cores underwent post-acquisition processing by linear spectral unmixing and deconvolved using the InForm software (version 3.1, Akoya Biosciences), and the tiles were stitched using Halo (version 3.6, Indica Labs). Tissue segmentation of the images was performed using a deep learning classifier by training the algorithm “DenseNet V2” from the Halo AI plug-in (version 3.6, Indica Labs), using only the DAPI channel as information for the training. Eight different classes were defined for both tissue segmentation and quality control issues: background, stroma, malignant tumor, benign glands, necrosis, liver tissue, decalcified tissue, and out of focus regions. The performance of the classifier was evaluated by a pathologist (F.S.) to ensure that the great majority of tissue compartments were properly classified. The cell segmentation was performed using a pre-trained deep learning model already present in Halo AI (“nuclei seg”) and applied to the DAPI channel only. Using the module “analysis” in the Halo software, all the biomarkers were then quantified using the segmentation algorithms to generate a counts matrix representing the average expression of each marker in each cell. Thresholding for each marker was performed using a gaussian mixture model of statistically robust cut-off values for low versus high intensity of the markers.

The *x* and *y*-coordinates for the precise nuclei or cytoplasmic marker locations from the immunofluorescence intensities were mapped for each TMA core. The coordinate point location was taken for multiple-localization analysis in different classes of markers. There are Mn classes of markers where each class is composed of a different combination of markers. For any given combination of markers, M1, …, Mn (each composed of nuclei or cytoplasm as detected with thresholds on their intensities), we classified a given nucleus or cytoplasm NCi coordinate, k in [1, …, i], belonging to M1, …, Mn class. We repeated this process for each NCi in M1, …, Mn to form a distribution of multiple-localizations in the different classes of markers, Ci. Measures for Ci were recorded for each TMA core sample and normalized using the total number of NCi belonging to M1, …, Mn class per patient. For any given marker, M1, …, Mn, we also measured the overall distribution at the single-localization level. The mean value for the measures was implemented by Welch ANOVA or unpaired t test (two-tailed) in Prism Version 9.5.0.

### Flow sorting of transfected cells

For CRISPR/Cas9-mediated gene knockout experiments, organoids were transfected with the lentiCas9-blast plasmid and blasticidin selection to establish stable lines. A custom vector for the sgRNA lentivirus carrying a TurboRFP reporter and puromycin antibiotic was then transfected into the Cas9-expressing organoids. After 14 days of antibiotic selection, we performed fluorescence-activated cell sorting (FACS) on the PE channel to sort RFP-positive (RFP^+^) cells on an BD Influx™ cell sorter, as described above. Approximately 1 x 10^6^ RFP^+^ cells were collected in 0.04% BSA in PBS (Miltenyi Biotec 130-091-376) for single-nuclei isolation and multi-ome ATAC/sequencing (10X Genomics). Additional collected RFP^+^ cells were used for organoid culture.

For experiments in which mouse NE tumor organoids were transfected with a HA-tagged H3.3K36M lentiviral vector, we performed flow cytometry at 14 days after antibiotic selection to determine the purity of the culture. Organoids were dissociated into single cells, and resuspended in 100 µl 4% paraformaldehyde/1 x 10^6^ cells to fix for 15 min at room temperature (RT). The fixed cells were neutralized with 1 ml PBS, washed once with PBS, and resuspended in 0.5 ml PBS. The cells were permeabilized for 10 min by addition of 0.5 ml 1% Triton-X100 with gentle vortexing to a final concentration of 0.5% Triton-X100. Cells were washed in 10 ml PBS, and resuspended with 100 µl Cy5.5® Conjugated mouse anti-HA-Tag antibody (Clone 6E2, Cell Signaling Technology 62145) diluted 1:50 in 0.5% BSA in PBS. Cells were incubated with antibody for 1 hour in the dark at RT, washed twice in 0.5% BSA, resuspended in 300 µl 0.5% BSA, and filtered through a Falcon™ Tube with 35 µm cell strainer cap. Flow cytometry was performed on the APC channel using a FACSCanto II Flow Cytometer (BD Bioscience) as described above, using Cy5.5 positive events to determine the percentage of HA-Tag^+^ cells. Flow data were analyzed using FlowJo (BD, version 10.8.2). The same batch of cells was collected in 0.04% BSA in PBS for single nuclei isolation and multiome snATAC/snRNA-sequencing.

### Single-nuclei multiome ATAC/RNA-sequencing

Organoids were digested into single cells using TrypLE, passed through a 40 µM cell strainer three times, and cell numbers were quantified using an automated cell counter. Approximately 1 x 10^6^ cells in 0.04% BSA/PBS (Miltenyi Biotec 130-091-376) were used for single nuclei isolation. Cells were spun at 1000 rpm for 5 min at 4°C, dissociated for 5 min with 100 µl ice-cold lysis buffer, neutralized with 1 ml wash buffer, spun at 1000 rpm for 5 min at 4°C, and resuspended in chilled Nuclei buffer included in the Single Cell Multiome ATAC Kit (10X Genomics). Wash buffer contained 10 mM Tris-HCl (pH 7.4), 10 mM NaCl, 3 mM MgCl_2_, 1% BAS, 0.1% Tween-20, 1 mM DTT (MilliporeSigma 646563), 1 U/µl RNase inhibitor (MilliporeSigma 3335399001) in nuclease-free water. Lysis buffer contained 10 mM Tris-HCl (pH 7.4), 10 mM NaCl, 3 mM MgCl_2_, 1% BAS, 0.1% Tween-20, 0.1% Nonidet P40 Substitute, 0.01% Digitonin, 1 mM DTT, 1 U/µl RNase inhibitor in nuclease-free water. Nuclei concentration was determined using a Countess II FL Automated Cell Counter.

Approximately 5,000 nuclei were loaded onto the Chromium Controller at the Columbia University Single Cell Analysis Core. Single-cell multiome ATAC/RNA-seq libraries were prepared following the manufacturer’s instructions (Chromium Next GEM Single Cell Multiome Reagent Kit, 10X Genomics). Briefly, nuclei suspensions were transposed and adapters added to the ends of the DNA fragments. Single Cell Multiome ATAC + GEX Gel Beads include a poly(dT) sequence that enables production of barcoded, full-length cDNA from poly-adenylated mRNA for the gene expression (GEX) library and a spacer sequence that enables barcode attachment to transposed DNA fragments for the ATAC library. The GEMs were generated by combining barcoded gel beads, transposed nuclei, and a master mix. Barcoded transposed DNA and barcoded full-length cDNA from poly-adenylated mRNA were amplified via PCR. Single-cell multiome ATAC/RNA-seq libraries were prepared for paired-end sequencing, and data were processed with Cell Ranger ARC software (10x Genomics, version 2.02) by the Columbia University Single Cell Analysis Core.

### Drug treatment of organoids

To generate drug response curves, organoids were digested with TrypLE for 10 min at 37°C, neutralized with PBS, gently dissociated into single cells, and passed through a 100 µm cell strainer. Cells were resuspended in 5% Matrigel in NE organoid culture medium lacking DHT and plated in triplicate at a seeding density of 5,000 cells/well in 96-well ultra-low attachment microplates. The next day, 7 doses of enzalutamide in 0.1% DMSO were dispensed at 1.5-fold dilution from 1 µM to 11.25 µM. Cell viability was assayed after five days using CellTiter-Glo 3D (Promega G9683), with luminescence was measured by a GloMax® Explorer multimode plate reader (Promega). Background luminescence was measured in medium without cells. The percentage of viable cells was calculated by the formula:

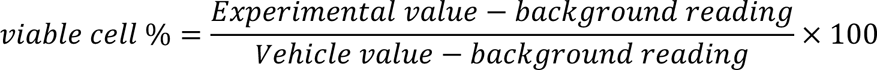

Drug response curves were generated by nonlinear regression using the percentage of viable cells against logarithm of drug concentrations using Graphpad Prism 9. IC_50_ values were calculated by the equation log(inhibitor) versus response (variable slope, four parameters). Two-way ANOVA was used to compare dose-response curves.

Similar methods were used to determine response of mouse or human organoids to defined doses of enzalutamide, using 5,000 cells/well (mouse) or 10,000 cells/well (human). The percentage of viable cells from different treatment groups were graphed using Prism 9. Unpaired t tests were used to compare means between two groups. All experiments were repeated independently at least three times with consistent results observed.

### Grafting assays

To generate tumors *in vivo*, mouse *NPp53* or human prostate organoids were grafted into 6-8 week old NOD/SCID male mice (NOD.CB17-*Prkdc^scid^/J*, Jackson Laboratory 001303). NOD/SCID mice underwent surgical castration at 7 days prior to grafting. For mouse grafts, 1 x 10^6^ dissociated organoid cells in 100 µl hepatocyte culture medium and 5% Matrigel were injected subcutaneously into the flank using a 1 ml syringe with 25G needle (BD 305122). For human grafts, 3 x 10^6^ dissociated cells in 100 µl 60% Matrigel and 40% hepatocyte culture medium were injected. Tumor sizes were measured with a digital caliper. In each cohort, 20-24 mice whose tumor volume had reached ∼250 mm^3^ (mouse grafts) or ∼80 mm^3^ (human grafts) at week two after grafting received either 10 mg/kg enzalutamide (TargetMol T6002) or 0.5% DMSO (MilliporeSigma D2650) by daily gavage via 20G needle (Roboz FN-7910) for 14 days (mouse grafts) or 56 days (human grafts). Enzalutamide or DMSO was suspended in 1% carboxymethylcellulose (MilliporeSigma 419281) and 0.1% Tween 80 (MilliporeSigma P4780) in distilled water. At the end of drug treatment, tumors were harvested and imaged under a stereo microscope (Olympus SZX16 with DP71 digital camera) using Olympus DP Controller 3.3.1.292 (Olympus). Tumor tissues were fixed in 10% formalin for 24-48 h and processed in the Columbia Molecular Pathology Core Facility. Tumor volumes were calculated using the formula:

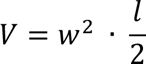

Tumor growth curves were plotted using Prism 9. Unpaired *t* tests were used to compare means between two groups, and *P* values were from two-tailed *t* tests.

### Single-cell and single-nuclei RNA-seq pre-processing

Single-cell RNA-sequencing (scRNA-seq) and single-nuclei RNA-sequencing (snRNA-seq) samples were processed independently using scanpy v1.9.1. for Python 3.9^72^. Cells with more than 1,000 detected genes and 1,000 UMI counts were retained, whereas cells with more than 10,000 detected genes or 50,000 UMI counts and with >20% of mitochondrial gene content were discarded. For doublet detection and removal, we used the Scrublet algorithm as implemented in scanpy, applied to each sample independently. Each sample was then subsampled by retaining 3,000 cells that passed quality control, using the *subsample* function with random state set to 666 as implemented in scanpy. All samples processed with the same technology (*i.e.,* single-cell or single-nuclei) were merged and UMI counts per cell were converted to sum to 1e4 and log-normalized.

### Reverse-engineering of prostate organoid regulatory networks

For each scRNA-seq sample, a Shared Neighbors Graph (SSN) was built with knn=15 to select cells with the most similar transcriptional profiles and merge them to generate high resolution ensembles of cells (metacells). This approach augments the number of detected genes per cell, which usually is very low due to technological dropout bias (<20%), thus increasing the number of targets that can be recovered by reverse-engineering of regulatory networks. Metacell profiles were computed on normalized data, merged into UMI counts, and transformed to count per million (cpm) for downstream analysis.

A sample-specific regulatory network (interactome) was reverse engineered from the resulting metacell cpm profiles (n = 500) using ARACNe-AP^41^, the most recent implementation of the ARACNe algorithm^40^, with 200 bootstraps, a Mutual Information (MI) P-value threshold P ≤ 10-8, and Data Processing Inequality (DPI) enabled. Regulatory proteins (RP) were selected into manually curated protein sets, including Transcription Factors (TF), co-Transcription Factors or chromatin remodeling enzymes, using the Gene Ontology (GO) identifiers GO:0003700 and GO:0003712. Each RP regulon (RP gene targets) was integrated across all the reverse-engineered networks to realize one final network. To avoid bias due to different regulon sizes, regulons were pruned to include only the 50 highest likelihood targets^39^, and regulons with <50 targets were excluded from the analysis. Mouse prostate cancer scRNA-seq and snRNA-seq gene expression profiles were scaled independently and transformed to protein activity profiles using the metacell-derived regulatory network and the VIPER algorithm. All samples were merged together to be projected on a 2-dimentional pane using Diffusion Maps^73^.

To recover cell identities, clusters of cells that share the same regulatory programs were identified using the Leiden clustering algorithm applied to all of the cells. Using these profiles, a Nearest Neighbor graph was built with knn=51 using the first 10 Principal Components to capture most of the variance. We performed a grid search analysis to tune the Leiden resolution parameter to maximize the average of within-cluster silhouette scores across each candidate optimal clustering solution. A high silhouette score is an indication that clustered cells have homogenous profiles, as sampled from the same cell population. The optimal solution yielded 3 major clusters.

### Single-nuclei ATAC-seq processing and analysis

To analyze snATAC-seq profiles, we used Signac v1.9.0 for R v4.1.2^74^. Peak sets were collected independently for each sample, as generated by the standard 10x Genomics Cell Ranger peak calling pipeline for scATAC-seq. To compare samples with distinct peak sets, a common set of genomic regions across all the samples was identified using GenomicRange v1.46. Genomic regions were annotated using UCSC annotation for mm10 as present in the EnsDb.Mmusculus.v79 database. Samples were integrated using the *FindIntegrationAnchors* function from Seurat v4.3.0 and the standard pipeline was then followed. Canonical AR binding sites were identified using the AR canonical motif (ID=MA0007.3) as present in JASPAR2020. AR inhibitor sensitive and resistant cells were defined based on sample conditions and VIPER-based clustering analysis (selecting cells from Cluster 1 across conditions). Differential peaks were shown using the *RegionHeatmap* function in genomic windows of 1,000 bp upstream and downstream of identified AR binding sites.

### Integration of CUT&Tag and VIPER analysis of snRNA-seq data

Histone mark count matrices were processed using limma *voom* (v3.54.2). Linear models were fitted for each gene based on the *voom* transformed data, and moderated t-statistics computed using the *eBayes* method from limma. A contrast matrix was built to compare differential H3K36me2 histone marks between untreated cells and NSD2 KO or H3.3K36M transfection. Top-ranked genes were extracted using the *topTable* function and genes with log-fold change < -0.5 and FDR < 0.1 were retained. This gene set was then used for GSEA analysis on gene expression signatures computed as differential between Cluster 1 and Cluster 3 across conditions (*e.g.,* NPPO-1NE-sgNSD2 Cluster 1 vs NPPO-1NE sgCtrl Cluster 3; Extended Data Fig. 7). For differential gene expression analysis Seurat v4.1.3 was used, with parameter *test.use* set to “DESeq2” in the *FindMarkers* function.

### Data availability

Single-cell RNA-sequencing and ATAC-sequencing data as well as CUT&Tag and bulk RNA-sequencing data from this study have been deposited in the Gene Expression Omnibus (GEO) under the accession number GSE237197.

## References

1. Watson, P.A., Arora, V.K. & Sawyers, C.L. Emerging mechanisms of resistance to androgen receptor inhibitors in prostate cancer. Nat Rev Cancer 15, 701–711 (2015).

2. Beltran, H., Hruszkewycz, A., Scher, H.I., Hildesheim, J., Isaacs, J., Yu, E.Y., Kelly, K., Lin, D., Dicker, A., Arnold, J., Hecht, T., Wicha, M., Sears, R., Rowley, D., White, R., Gulley, J.L., Lee, J., Diaz Meco, M., Small, E.J., Shen, M., Knudsen, K., Goodrich, D.W., Lotan, T., Zoubeidi, A., Sawyers, C.L., Rudin, C.M., Loda, M., Thompson, T., Rubin, M.A., Tawab-Amiri, A., Dahut, W. & Nelson, P.S. The role of lineage plasticity in prostate cancer therapy resistance. Clin Cancer Res 25, 6916–6924 (2019).

3. Davies, A.H., Beltran, H. & Zoubeidi, A. Cellular plasticity and the neuroendocrine phenotype in prostate cancer. Nat Rev Urol 15, 271–286 (2018).

4. Rickman, D.S., Beltran, H., Demichelis, F. & Rubin, M.A. Biology and evolution of poorly differentiated neuroendocrine tumors. Nat Med 23, 1–10 (2017).

5. Rubin, M.A., Bristow, R.G., Thienger, P.D., Dive, C. & Imielinski, M. Impact of lineage plasticity to and from a neuroendocrine phenotype on progression and response in prostate and lung cancers. Mol Cell 80, 562–577 (2020).

6. Bennett, R.L., Swaroop, A., Troche, C. & Licht, J.D. The role of nuclear receptor-binding SET domain family histone lysine methyltransferases in cancer. Cold Spring Harb Perspect Med 7, a026708 (2017).

7. Zou, M., Toivanen, R., Mitrofanova, A., Floch, N., Hayati, S., Sun, Y., Le Magnen, C., Chester, D., Mostaghel, E.A., Califano, A., Rubin, M.A., Shen, M.M. & Abate-Shen, C. Transdifferentiation as a mechanism of treatment resistance in a mouse model of castration-resistant prostate cancer. Cancer Discov 7, 736–749 (2017).

8. Hanahan, D. Hallmarks of cancer: new dimensions. Cancer Discov 12, 31–46 (2022).

9. Beltran, H., Rickman, D.S., Park, K., Chae, S.S., Sboner, A., MacDonald, T.Y., Wang, Y., Sheikh, K.L., Terry, S., Tagawa, S.T., Dhir, R., Nelson, J.B., de la Taille, A., Allory, Y., Gerstein, M.B., Perner, S., Pienta, K.J., Chinnaiyan, A.M., Wang, Y., Collins, C.C., Gleave, M.E., Demichelis, F., Nanus, D.M. & Rubin, M.A. Molecular characterization of neuroendocrine prostate cancer and identification of new drug targets. Cancer Discov 1, 487–495 (2011).

10. Quigley, D.A., Dang, H.X., Zhao, S.G., Lloyd, P., Aggarwal, R., Alumkal, J.J., Foye, A., Kothari, V., Perry, M.D., Bailey, A.M., Playdle, D., Barnard, T.J., Zhang, L., Zhang, J., Youngren, J.F., Cieslik, M.P., Parolia, A., Beer, T.M., Thomas, G., Chi, K.N., Gleave, M., Lack, N.A., Zoubeidi, A., Reiter, R.E., Rettig, M.B., Witte, O., Ryan, C.J., Fong, L., Kim, W., Friedlander, T., Chou, J., Li, H., Das, R., Li, H., Moussavi-Baygi, R., Goodarzi, H., Gilbert, L.A., Lara, P.N., Jr., Evans, C.P., Goldstein, T.C., Stuart, J.M., Tomlins, S.A., Spratt, D.E., Cheetham, R.K., Cheng, D.T., Farh, K., Gehring, J.S., Hakenberg, J., Liao, A., Febbo, P.G., Shon, J., Sickler, B., Batzoglou, S., Knudsen, K.E., He, H.H., Huang, J., Wyatt, A.W., Dehm, S.M., Ashworth, A., Chinnaiyan, A.M., Maher, C.A., Small, E.J. & Feng, F.Y. Genomic hallmarks and structural variation in metastatic prostate cancer. Cell 174, 758–769 (2018).

11. Le Magnen, C., Shen, M.M. & Abate-Shen, C. Lineage plasticity in cancer progression and treatment. Annu Rev Cancer Biol 2, 271–289 (2018).

12. Labrecque, M.P., Coleman, I.M., Brown, L.G., True, L.D., Kollath, L., Lakely, B., Nguyen, H.M., Yang, Y.C., da Costa, R.M.G., Kaipainen, A., Coleman, R., Higano, C.S., Yu, E.Y., Cheng, H.H., Mostaghel, E.A., Montgomery, B., Schweizer, M.T., Hsieh, A.C., Lin, D.W., Corey, E., Nelson, P.S. & Morrissey, C. Molecular profiling stratifies diverse phenotypes of treatment-refractory metastatic castration-resistant prostate cancer. J Clin Invest 130, 4492–4505 (2019).

13. Bluemn, E.G., Coleman, I.M., Lucas, J.M., Coleman, R.T., Hernandez-Lopez, S., Tharakan, R., Bianchi-Frias, D., Dumpit, R.F., Kaipainen, A., Corella, A.N., Yang, Y.C., Nyquist, M.D., Mostaghel, E., Hsieh, A.C., Zhang, X., Corey, E., Brown, L.G., Nguyen, H.M., Pienta, K., Ittmann, M., Schweizer, M., True, L.D., Wise, D., Rennie, P.S., Vessella, R.L., Morrissey, C. & Nelson, P.S. Androgen receptor pathway-independent prostate cancer is sustained through FGF signaling. Cancer Cell 32, 474–489 (2017).

14. Su, W., Han, H.H., Wang, Y., Zhang, B., Zhou, B., Cheng, Y.K., Rumandia, A., Gurrapu, S., Chakraborty, G., Su, J., Yang, G., Liang, X., Wang, G., Rosen, N., Scher, H.I., Ouerfelli, O. & Giancotti, F. The Polycomb Repressor Complex 1 drives double negative prostate cancer metastasis by coordinating stemness and immune suppression. Cancer Cell 36, 139–155 (2019).

15. Zaffuto, E., Pompe, R., Zanaty, M., Bondarenko, H.D., Leyh-Bannurah, S.R., Moschini, M., Dell’Oglio, P., Gandaglia, G., Fossati, N., Stabile, A., Zorn, K.C., Montorsi, F., Briganti, A. & Karakiewicz, P.I. Contemporary incidence and cancer control outcomes of primary neuroendocrine prostate cancer: A SEER database analysis. Clin Genitourin Cancer 15, e793–e800 (2017).

16. Aggarwal, R., Huang, J., Alumkal, J.J., Zhang, L., Feng, F.Y., Thomas, G.V., Weinstein, A.S., Friedl, V., Zhang, C., Witte, O.N., Lloyd, P., Gleave, M., Evans, C.P., Youngren, J., Beer, T.M., Rettig, M., Wong, C.K., True, L., Foye, A., Playdle, D., Ryan, C.J., Lara, P., Chi, K.N., Uzunangelov, V., Sokolov, A., Newton, Y., Beltran, H., Demichelis, F., Rubin, M.A., Stuart, J.M. & Small, E.J. Clinical and genomic characterization of treatment-emergent small-cell neuroendocrine prostate cancer: a multi-institutional prospective study. J Clin Oncol 36, 2492–2503 (2018).

17. Abida, W., Cyrta, J., Heller, G., Prandi, D., Armenia, J., Coleman, I., Cieslik, M., Benelli, M., Robinson, D., Van Allen, E.M., Sboner, A., Fedrizzi, T., Mosquera, J.M., Robinson, B.D., De Sarkar, N., Kunju, L.P., Tomlins, S., Wu, Y.M., Nava Rodrigues, D., Loda, M., Gopalan, A., Reuter, V.E., Pritchard, C.C., Mateo, J., Bianchini, D., Miranda, S., Carreira, S., Rescigno, P., Filipenko, J., Vinson, J., Montgomery, R.B., Beltran, H., Heath, E.I., Scher, H.I., Kantoff, P.W., Taplin, M.E., Schultz, N., deBono, J.S., Demichelis, F., Nelson, P.S., Rubin, M.A., Chinnaiyan, A.M. & Sawyers, C.L. Genomic correlates of clinical outcome in advanced prostate cancer. Proc Natl Acad Sci USA 116, 11428–11436 (2019).

18. Laudato, S., Aparicio, A. & Giancotti, F.G. Clonal evolution and epithelial plasticity in the emergence of AR-independent prostate carcinoma. Trends Cancer 5, 440–455 (2019).

19. Tang, D.G. Understanding and targeting prostate cancer cell heterogeneity and plasticity. Semin Cancer Biol 82, 68–93 (2022).

20. Li, W. & Shen, M.M. Prostate cancer cell heterogeneity and plasticity: Insights from studies of genetically-engineered mouse models. Semin Cancer Biol 82, 60–67 (2022).

21. Quintanal-Villalonga, A., Chan, J.M., Yu, H.A., Pe’er, D., Sawyers, C.L., Sen, T. & Rudin, C.M. Lineage plasticity in cancer: a shared pathway of therapeutic resistance. Nat Rev Clin Oncol 17, 360–371 (2020).

22. Chan, J.M., Zaidi, S., Love, J.R., Zhao, J.L., Setty, M., Wadosky, K.M., Gopalan, A., Choo, Z.N., Persad, S., Choi, J., LaClair, J., Lawrence, K.E., Chaudhary, O., Xu, T., Masilionis, I., Linkov, I., Wang, S., Lee, C., Barlas, A., Morris, M.J., Mazutis, L., Chaligne, R., Chen, Y., Goodrich, D.W., Karthaus, W.R., Pe’er, D. & Sawyers, C.L. Lineage plasticity in prostate cancer depends on JAK/STAT inflammatory signaling. Science 377, 1180–1191 (2022).

23. Beltran, H., Prandi, D., Mosquera, J.M., Benelli, M., Puca, L., Cyrta, J., Marotz, C., Giannopoulou, E., Chakravarthi, B.V., Varambally, S., Tomlins, S.A., Nanus, D.M., Tagawa, S.T., Van Allen, E.M., Elemento, O., Sboner, A., Garraway, L.A., Rubin, M.A. & Demichelis, F. Divergent clonal evolution of castration-resistant neuroendocrine prostate cancer. Nat Med 22, 298–305 (2016).

24. Varambally, S., Dhanasekaran, S.M., Zhou, M., Barrette, T.R., Kumar-Sinha, C., Sanda, M.G., Ghosh, D., Pienta, K.J., Sewalt, R.G., Otte, A.P., Rubin, M.A. & Chinnaiyan, A.M. The polycomb group protein EZH2 is involved in progression of prostate cancer. Nature 419, 624–629 (2002).

25. Dardenne, E., Beltran, H., Benelli, M., Gayvert, K., Berger, A., Puca, L., Cyrta, J., Sboner, A., Noorzad, Z., MacDonald, T., Cheung, C., Yuen, K.S., Gao, D., Chen, Y., Eilers, M., Mosquera, J.M., Robinson, B.D., Elemento, O., Rubin, M.A., Demichelis, F. & Rickman, D.S. N-Myc induces an EZH2-mediated transcriptional program driving neuroendocrine prostate cancer. Cancer Cell 30, 563–577 (2016).

26. Ku, S.Y., Rosario, S., Wang, Y., Mu, P., Seshadri, M., Goodrich, Z.W., Goodrich, M.M., Labbe, D.P., Gomez, E.C., Wang, J., Long, H.W., Xu, B., Brown, M., Loda, M., Sawyers, C.L., Ellis, L. & Goodrich, D.W. Rb1 and Trp53 cooperate to suppress prostate cancer lineage plasticity, metastasis, and antiandrogen resistance. Science 355, 78–83 (2017).

27. Puca, L., Bareja, R., Prandi, D., Shaw, R., Benelli, M., Karthaus, W.R., Hess, J., Sigouros, M., Donoghue, A., Kossai, M., Gao, D., Cyrta, J., Sailer, V., Vosoughi, A., Pauli, C., Churakova, Y., Cheung, C., Deonarine, L.D., McNary, T.J., Rosati, R., Tagawa, S.T., Nanus, D.M., Mosquera, J.M., Sawyers, C.L., Chen, Y., Inghirami, G., Rao, R.A., Grandori, C., Elemento, O., Sboner, A., Demichelis, F., Rubin, M.A. & Beltran, H. Patient derived organoids to model rare prostate cancer phenotypes. Nat Commun 9, 2404 (2018).

28. Li, Y., Trojer, P., Xu, C.F., Cheung, P., Kuo, A., Drury, W.J., 3rd, Qiao, Q., Neubert, T.A., Xu, R.M., Gozani, O. & Reinberg, D. The target of the NSD family of histone lysine methyltransferases depends on the nature of the substrate. J Biol Chem 284, 34283–34295 (2009).

29. Kuo, A.J., Cheung, P., Chen, K., Zee, B.M., Kioi, M., Lauring, J., Xi, Y., Park, B.H., Shi, X., Garcia, B.A., Li, W. & Gozani, O. NSD2 links dimethylation of histone H3 at lysine 36 to oncogenic programming. Mol Cell 44, 609–620 (2011).

30. Martinez-Garcia, E., Popovic, R., Min, D.J., Sweet, S.M., Thomas, P.M., Zamdborg, L., Heffner, A., Will, C., Lamy, L., Staudt, L.M., Levens, D.L., Kelleher, N.L. & Licht, J.D. The MMSET histone methyl transferase switches global histone methylation and alters gene expression in t(4;14) multiple myeloma cells. Blood 117, 211–220 (2011).

31. Popovic, R., Martinez-Garcia, E., Giannopoulou, E.G., Zhang, Q., Zhang, Q., Ezponda, T., Shah, M.Y., Zheng, Y., Will, C.M., Small, E.C., Hua, Y., Bulic, M., Jiang, Y., Carrara, M., Calogero, R.A., Kath, W.L., Kelleher, N.L., Wang, J.P., Elemento, O. & Licht, J.D. Histone methyltransferase MMSET/NSD2 alters EZH2 binding and reprograms the myeloma epigenome through global and focal changes in H3K36 and H3K27 methylation. PLoS Genet 10, e1004566 (2014).

32. Streubel, G., Watson, A., Jammula, S.G., Scelfo, A., Fitzpatrick, D.J., Oliviero, G., McCole, R., Conway, E., Glancy, E., Negri, G.L., Dillon, E., Wynne, K., Pasini, D., Krogan, N.J., Bracken, A.P. & Cagney, G. The H3K36me2 methyltransferase Nsd1 demarcates PRC2-mediated H3K27me2 and H3K27me3 domains in embryonic stem cells. Mol Cell 70, 371–379 e375 (2018).

33. Yuan, W., Xu, M., Huang, C., Liu, N., Chen, S. & Zhu, B. H3K36 methylation antagonizes PRC2-mediated H3K27 methylation. J Biol Chem 286, 7983–7989 (2011).

34. Schmitges, F.W., Prusty, A.B., Faty, M., Stutzer, A., Lingaraju, G.M., Aiwazian, J., Sack, R., Hess, D., Li, L., Zhou, S., Bunker, R.D., Wirth, U., Bouwmeester, T., Bauer, A., Ly-Hartig, N., Zhao, K., Chan, H., Gu, J., Gut, H., Fischle, W., Muller, J. & Thoma, N.H. Histone methylation by PRC2 is inhibited by active chromatin marks. Mol Cell 42, 330–341 (2011).

35. Weinberg, D.N., Papillon-Cavanagh, S., Chen, H., Yue, Y., Chen, X., Rajagopalan, K.N., Horth, C., McGuire, J.T., Xu, X., Nikbakht, H., Lemiesz, A.E., Marchione, D.M., Marunde, M.R., Meiners, M.J., Cheek, M.A., Keogh, M.C., Bareke, E., Djedid, A., Harutyunyan, A.S., Jabado, N., Garcia, B.A., Li, H., Allis, C.D., Majewski, J. & Lu, C. The histone mark H3K36me2 recruits DNMT3A and shapes the intergenic DNA methylation landscape. Nature 573, 281–286 (2019).

36. Shirane, K., Miura, F., Ito, T. & Lorincz, M.C. NSD1-deposited H3K36me2 directs de novo methylation in the mouse male germline and counteracts Polycomb-associated silencing. Nat Genet 52, 1088–1098 (2020).

37. Hamagami, N., Wu, D.Y., Clemens, A.W., Nettles, S.A., Li, A. & Gabel, H.W. NSD1 deposits histone H3 lysine 36 dimethylation to pattern non-CG DNA methylation in neurons. Mol Cell 83, 1412–1428 (2023).

38. Wang, X., Kruithof-de Julio, M., Economides, K.D., Walker, D., Yu, H.L., Halili, M.V., Hu, Y.P., Price, S.M., Abate-Shen, C. & Shen, M.M. A luminal epithelial stem cell that is a cell of origin for prostate cancer. Nature 461, 495–500 (2009).

39. Alvarez, M.J., Shen, Y., Giorgi, F.M., Lachmann, A., Ding, B.B., Ye, B.H. & Califano, A. Functional characterization of somatic mutations in cancer using network-based inference of protein activity. Nat Genet 48, 838–847 (2016).

40. Margolin, A.A., Nemenman, I., Basso, K., Wiggins, C., Stolovitzky, G., Dalla Favera, R. & Califano, A. ARACNE: an algorithm for the reconstruction of gene regulatory networks in a mammalian cellular context. BMC Bioinformatics 7 Suppl 1, S7 (2006).

41. Lachmann, A., Giorgi, F.M., Lopez, G. & Califano, A. ARACNe-AP: gene network reverse engineering through adaptive partitioning inference of mutual information. Bioinformatics 32, 2233–2235 (2016).

42. Gulati, G.S., Sikandar, S.S., Wesche, D.J., Manjunath, A., Bharadwaj, A., Berger, M.J., Ilagan, F., Kuo, A.H., Hsieh, R.W., Cai, S., Zabala, M., Scheeren, F.A., Lobo, N.A., Qian, D., Yu, F.B., Dirbas, F.M., Clarke, M.F. & Newman, A.M. Single-cell transcriptional diversity is a hallmark of developmental potential. Science 367, 405–411 (2020).

43. Han, H., Wang, Y., Curto, J., Gurrapu, S., Laudato, S., Rumandla, A., Chakraborty, G., Wang, X., Chen, H., Jiang, Y., Kumar, D., Caggiano, E.G., Capogiri, M., Zhang, B., Ji, Y., Maity, S.N., Hu, M., Bai, S., Aparicio, A.M., Efstathiou, E., Logothetis, C.J., Navin, N., Navone, N.M., Chen, Y. & Giancotti, F.G. Mesenchymal and stem-like prostate cancer linked to therapy-induced lineage plasticity and metastasis. Cell Rep 39, 110595 (2022).

44. Kaya-Okur, H.S., Wu, S.J., Codomo, C.A., Pledger, E.S., Bryson, T.D., Henikoff, J.G., Ahmad, K. & Henikoff, S. CUT&Tag for efficient epigenomic profiling of small samples and single cells. Nat Commun 10, 1930 (2019).

45. Sun, Z., Lin, Y., Islam, M.T., Koche, R., Hedehus, L., Liu, D., Huang, C., Vierbuchen, T., Sawyers, C.L. & Helin, K. Chromatin regulation of transcriptional enhancers and cell fate by the Sotos syndrome gene NSD1. Mol Cell, doi: 10.1016/j.molcel.2023.1006.1007 (2023).

46. Aytes, A., Giacobbe, A., Mitrofanova, A., Ruggero, K., Cyrta, J., Arriaga, J., Palomero, L., Farran-Matas, S., Rubin, M.A., Shen, M.M., Califano, A. & Abate-Shen, C. NSD2 is a conserved driver of metastatic prostate cancer progression. Nat Commun 9, 5201 (2018).

47. Yang, P., Guo, L., Duan, Z.J., Tepper, C.G., Xue, L., Chen, X., Kung, H.J., Gao, A.C., Zou, J.X. & Chen, H.W. Histone methyltransferase NSD2/MMSET mediates constitutive NF-kappaB signaling for cancer cell proliferation, survival, and tumor growth via a feed-forward loop. Mol Cell Biol 32, 3121–3131 (2012).

48. Filon, M., Gawdzik, J., Truong, A., Allen, G., Huang, W., Khemees, T., Machhi, R., Lewis, P., Yang, B., Denu, J. & Jarrard, D. Tandem histone methyltransferase upregulation defines a unique aggressive prostate cancer phenotype. Br J Cancer 125, 247–254 (2021).

49. Lu, C., Jain, S.U., Hoelper, D., Bechet, D., Molden, R.C., Ran, L., Murphy, D., Venneti, S., Hameed, M., Pawel, B.R., Wunder, J.S., Dickson, B.C., Lundgren, S.M., Jani, K.S., De Jay, N., Papillon-Cavanagh, S., Andrulis, I.L., Sawyer, S.L., Grynspan, D., Turcotte, R.E., Nadaf, J., Fahiminiyah, S., Muir, T.W., Majewski, J., Thompson, C.B., Chi, P., Garcia, B.A., Allis, C.D., Jabado, N. & Lewis, P.W. Histone H3K36 mutations promote sarcomagenesis through altered histone methylation landscape. Science 352, 844–849 (2016).

50. Fang, D., Gan, H., Lee, J.H., Han, J., Wang, Z., Riester, S.M., Jin, L., Chen, J., Zhou, H., Wang, J., Zhang, H., Yang, N., Bradley, E.W., Ho, T.H., Rubin, B.P., Bridge, J.A., Thibodeau, S.N., Ordog, T., Chen, Y., van Wijnen, A.J., Oliveira, A.M., Xu, R.M., Westendorf, J.J. & Zhang, Z. The histone H3.3K36M mutation reprograms the epigenome of chondroblastomas. Science 352, 1344–1348 (2016).

51. Rajagopalan, K.N., Chen, X., Weinberg, D.N., Chen, H., Majewski, J., Allis, C.D. & Lu, C. Depletion of H3K36me2 recapitulates epigenomic and phenotypic changes induced by the H3.3K36M oncohistone mutation. Proc Natl Acad Sci USA 118, e2021795118 (2021).

52. Tang, F., Xu, D., Wang, S., Wong, C.K., Martinez-Fundichely, A., Lee, C.J., Cohen, S., Park, J., Hill, C.E., Eng, K., Bareja, R., Han, T., Liu, E.M., Palladino, A., Di, W., Gao, D., Abida, W., Beg, S., Puca, L., Meneses, M., de Stanchina, E., Berger, M.F., Gopalan, A., Dow, L.E., Mosquera, J.M., Beltran, H., Sternberg, C.N., Chi, P., Scher, H.I., Sboner, A., Chen, Y. & Khurana, E. Chromatin profiles classify castration-resistant prostate cancers suggesting therapeutic targets. Science 376, eabe1505 (2022).

53. Pomerantz, M.M., Li, F., Takeda, D.Y., Lenci, R., Chonkar, A., Chabot, M., Cejas, P., Vazquez, F., Cook, J., Shivdasani, R.A., Bowden, M., Lis, R., Hahn, W.C., Kantoff, P.W., Brown, M., Loda, M., Long, H.W. & Freedman, M.L. The androgen receptor cistrome is extensively reprogrammed in human prostate tumorigenesis. Nat Genet 47, 1346–1351 (2015).

54. Pomerantz, M.M., Qiu, X., Zhu, Y., Takeda, D.Y., Pan, W., Baca, S.C., Gusev, A., Korthauer, K.D., Severson, T.M., Ha, G., Viswanathan, S.R., Seo, J.H., Nguyen, H.M., Zhang, B., Pasaniuc, B., Giambartolomei, C., Alaiwi, S.A., Bell, C.A., O’Connor, E.P., Chabot, M.S., Stillman, D.R., Lis, R., Font-Tello, A., Li, L., Cejas, P., Bergman, A.M., Sanders, J., van der Poel, H.G., Gayther, S.A., Lawrenson, K., Fonseca, M.A.S., Reddy, J., Corona, R.I., Martovetsky, G., Egan, B., Choueiri, T., Ellis, L., Garraway, I.P., Lee, G.M., Corey, E., Long, H.W., Zwart, W. & Freedman, M.L. Prostate cancer reactivates developmental epigenomic programs during metastatic progression. Nat Genet 52, 790–799 (2020).

55. Severson, T., Qiu, X., Alshalalfa, M., Sjostrom, M., Quigley, D., Bergman, A., Long, H., Feng, F., Freedman, M.L., Zwart, W. & Pomerantz, M.M. Androgen receptor reprogramming demarcates prognostic, context-dependent gene sets in primary and metastatic prostate cancer. Clin Epigenetics 14, 60 (2022).

56. Xiao, L., Parolia, A., Qiao, Y., Bawa, P., Eyunni, S., Mannan, R., Carson, S.E., Chang, Y., Wang, X., Zhang, Y., Vo, J.N., Kregel, S., Simko, S.A., Delekta, A.D., Jaber, M., Zheng, H., Apel, I.J., McMurry, L., Su, F., Wang, R., Zelenka-Wang, S., Sasmal, S., Khare, L., Mukherjee, S., Abbineni, C., Aithal, K., Bhakta, M.S., Ghurye, J., Cao, X., Navone, N.M., Nesvizhskii, A.I., Mehra, R., Vaishampayan, U., Blanchette, M., Wang, Y., Samajdar, S., Ramachandra, M. & Chinnaiyan, A.M. Targeting SWI/SNF ATPases in enhancer-addicted prostate cancer. Nature 601, 434–439 (2022).

57. Wang, Q., Li, W., Zhang, Y., Yuan, X., Xu, K., Yu, J., Chen, Z., Beroukhim, R., Wang, H., Lupien, M., Wu, T., Regan, M.M., Meyer, C.A., Carroll, J.S., Manrai, A.K., Janne, O.A., Balk, S.P., Mehra, R., Han, B., Chinnaiyan, A.M., Rubin, M.A., True, L., Fiorentino, M., Fiore, C., Loda, M., Kantoff, P.W., Liu, X.S. & Brown, M. Androgen receptor regulates a distinct transcription program in androgen-independent prostate cancer. Cell 138, 245–256 (2009).

58. Han, M., Li, F., Zhang, Y., Dai, P., He, J., Li, Y., Zhu, Y., Zheng, J., Huang, H., Bai, F. & Gao, D. FOXA2 drives lineage plasticity and KIT pathway activation in neuroendocrine prostate cancer. Cancer Cell 40, 1306–1323 (2022).

59. Rotinen, M., You, S., Yang, J., Coetzee, S.G., Reis-Sobreiro, M., Huang, W.C., Huang, F., Pan, X., Yanez, A., Hazelett, D.J., Chu, C.Y., Steadman, K., Morrissey, C.M., Nelson, P.S., Corey, E., Chung, L.W.K., Freedland, S.J., Di Vizio, D., Garraway, I.P., Murali, R., Knudsen, B.S. & Freeman, M.R. ONECUT2 is a targetable master regulator of lethal prostate cancer that suppresses the androgen axis. Nat Med 24, 1887–1898 (2018).

60. Kang, H.B., Choi, Y., Lee, J.M., Choi, K.C., Kim, H.C., Yoo, J.Y., Lee, Y.H. & Yoon, H.G. The histone methyltransferase, NSD2, enhances androgen receptor-mediated transcription. FEBS Lett 583, 1880–1886 (2009).

61. Li, N., Xue, W., Yuan, H., Dong, B., Ding, Y., Liu, Y., Jiang, M., Kan, S., Sun, T., Ren, J., Pan, Q., Li, X., Zhang, P., Hu, G., Wang, Y., Wang, X., Li, Q. & Qin, J. AKT-mediated stabilization of histone methyltransferase WHSC1 promotes prostate cancer metastasis. J Clin Invest 127, 1284–1302 (2017).

62. Ezponda, T., Popovic, R., Shah, M.Y., Martinez-Garcia, E., Zheng, Y., Min, D.J., Will, C., Neri, A., Kelleher, N.L., Yu, J. & Licht, J.D. The histone methyltransferase MMSET/WHSC1 activates TWIST1 to promote an epithelial-mesenchymal transition and invasive properties of prostate cancer. Oncogene 32, 2882–2890 (2013).

63. Yuan, S., Natesan, R., Sanchez-Rivera, F.J., Li, J., Bhanu, N.V., Yamazoe, T., Lin, J.H., Merrell, A.J., Sela, Y., Thomas, S.K., Jiang, Y., Plesset, J.B., Miller, E.M., Shi, J., Garcia, B.A., Lowe, S.W., Asangani, I.A. & Stanger, B.Z. Global regulation of the histone mark H3K36me2 underlies epithelial plasticity and metastatic progression. Cancer Discov 10, 854–871 (2020).

64. Chesi, M., Nardini, E., Lim, R.S., Smith, K.D., Kuehl, W.M. & Bergsagel, P.L. The t(4;14) translocation in myeloma dysregulates both FGFR3 and a novel gene, MMSET, resulting in IgH/MMSET hybrid transcripts. Blood 92, 3025–3034 (1998).

65. Jaffe, J.D., Wang, Y., Chan, H.M., Zhang, J., Huether, R., Kryukov, G.V., Bhang, H.E., Taylor, J.E., Hu, M., Englund, N.P., Yan, F., Wang, Z., Robert McDonald, E., 3rd, Wei, L., Ma, J., Easton, J., Yu, Z., deBeaumount, R., Gibaja, V., Venkatesan, K., Schlegel, R., Sellers, W.R., Keen, N., Liu, J., Caponigro, G., Barretina, J., Cooke, V.G., Mullighan, C., Carr, S.A., Downing, J.R., Garraway, L.A. & Stegmeier, F. Global chromatin profiling reveals NSD2 mutations in pediatric acute lymphoblastic leukemia. Nat Genet 45, 1386–1391 (2013).

66. Papillon-Cavanagh, S., Lu, C., Gayden, T., Mikael, L.G., Bechet, D., Karamboulas, C., Ailles, L., Karamchandani, J., Marchione, D.M., Garcia, B.A., Weinreb, I., Goldstein, D., Lewis, P.W., Dancu, O.M., Dhaliwal, S., Stecho, W., Howlett, C.J., Mymryk, J.S., Barrett, J.W., Nichols, A.C., Allis, C.D., Majewski, J. & Jabado, N. Impaired H3K36 methylation defines a subset of head and neck squamous cell carcinomas. Nat Genet 49, 180–185 (2017).

67. Yuan, G., Flores, N.M., Hausmann, S., Lofgren, S.M., Kharchenko, V., Angulo-Ibanez, M., Sengupta, D., Lu, X., Czaban, I., Azhibek, D., Vicent, S., Fischle, W., Jaremko, M., Fang, B., Wistuba, II, Chua, K.F., Roth, J.A., Minna, J.D., Shao, N.Y., Jaremko, L., Mazur, P.K. & Gozani, O. Elevated NSD3 histone methylation activity drives squamous cell lung cancer. Nature 590, 504–508 (2021).

68. Sengupta, D., Zeng, L., Li, Y., Hausmann, S., Ghosh, D., Yuan, G., Nguyen, T.N., Lyu, R., Caporicci, M., Morales Benitez, A., Coles, G.L., Kharchenko, V., Czaban, I., Azhibek, D., Fischle, W., Jaremko, M., Wistuba, II, Sage, J., Jaremko, L., Li, W., Mazur, P.K. & Gozani, O. NSD2 dimethylation at H3K36 promotes lung adenocarcinoma pathogenesis. Mol Cell 81, 4481–4492 e4489 (2021).

69. Wang, Z.A., Mitrofanova, A., Bergren, S.K., Abate-Shen, C., Cardiff, R.D., Califano, A. & Shen, M.M. Lineage analysis of basal epithelial cells reveals their unexpected plasticity and supports a cell-of-origin model for prostate cancer heterogeneity. Nat Cell Biol 15, 274–283 (2013).

70. Chua, C.W., Shibata, M., Lei, M., Toivanen, R., Barlow, L.J., Bergren, S.K., Badani, K.K., McKiernan, J.M., Benson, M.C., Hibshoosh, H. & Shen, M.M. Single luminal epithelial progenitors can generate prostate organoids in culture. Nat Cell Biol 16, 951–961 (2014).

71. Shechter, D., Dormann, H.L., Allis, C.D. & Hake, S.B. Extraction, purification and analysis of histones. Nat Protoc 2, 1445–1457 (2007).

72. Wolf, F.A., Angerer, P. & Theis, F.J. SCANPY: large-scale single-cell gene expression data analysis. Genome Biol 19, 15 (2018).

73. Haghverdi, L., Buettner, F. & Theis, F.J. Diffusion maps for high-dimensional single-cell analysis of differentiation data. Bioinformatics 31, 2989–2998 (2015).

74. Stuart, T., Srivastava, A., Madad, S., Lareau, C.A. & Satija, R. Single-cell chromatin state analysis with Signac. Nat Methods 18, 1333–1341 (2021).

